# Thermococcales pangenome uncovers new genera and modular evolution

**DOI:** 10.64898/2026.09.28.752058

**Authors:** Ryan Catchpole, Clémence Lauden, Damien Courtine, Evelyne Marguet, Stéphanie Fouteau, Catherine Badel, Michalis-Daniel Lazar, Vladislav Shutko, Ghislaine Magdelenat, Valérie Barbe, Jean Weissenbach, Anthony Woo, Philippe M Oger, Gaël Erauso, Loïs Maignien, Mohamed Jebbar, Patrick Forterre, Jacques Oberto, Violette Da Cunha

## Abstract

The order Thermococcales is unusual among Archaea in being known primarily from hundreds of lab-cultured isolates rather than metagenome-assembled genomes. Such cultivation is powerful for comparative genomics, allowing access to complete closed genomes rather than fragmented assemblies. We analysed 122 complete genomes, comprising all publicly available type species plus 67 new isolates, as a single pangenome. A combination of genome-scale metrics resolved 70 species across five genus-level clades, two proposed here as the new genera *Oleococcus* and *Piezococcus*, in addition to the canonical *Pyrococcus, Palaeococcus,* and *Thermococcus.* Only 6.1% of the 21,490 gene families are persistent, yet these contribute over half the genes in any genome. We observe that accessory functions, such as amino-acid biosynthesis, often behave modularly, being gained and lost as intact cassettes from genus-specific chromosomal loci. Combined with 114 mobile genetic elements, 72 integrated into the chromosome at conserved tRNA genes, and 42 extrachromosomal, we find that Thermococcales genomes accommodate rapid horizontal turnover through a constrained chromosomal architecture.

## Introduction

The order Thermococcales comprises anaerobic, sulfur-reducing, hyperthermophilic archaea that are among the most readily cultivated microorganisms of deep-sea hydrothermal vents, and as such have become workhorses of archaeal molecular biology. Members of the three described genera, *Pyrococcus, Thermococcus,* and *Palaeococcus* grow rapidly and to high density in defined media, tolerate a broad range of temperature and hydrostatic pressure, and support genetic systems in several species, including *T. kodakarensis, Py. furiosus, T. onnurineus, T. barophilus,* and *Py. yayanosii* [1–5]. Decades of work on their glycolytic and redox enzymes, DNA replication and repair machinery, and thermostable proteins have made Thermococcales model organisms for archaeal physiology and biotechnology [2,6,7]. Crucially, this experimental accessibility has been accompanied by a sustained isolation effort, with many hundreds of Thermococcales strains held in culture collections worldwide, isolated from hydrothermal vent fields across every major ocean basin as well as from terrestrial hot springs and subsurface oil reservoirs.

This cultured diversity makes Thermococcales an unusual case within archaeal biology, and especially archaeal genomics. The known diversity of Archaea has expanded rapidly in recent years through metagenome-assembled genomes, which have added dozens of candidate phyla but few cultured representatives [8–11]. Such MAGs are important for phylogenetic and metabolic inferences but limited the scope of research – phenotypes cannot be measured directly, genotypes cannot be perturbed, and formal naming requires isolated type strains [12–14]. Thermococcales invert this paradigm, with dense sampling of strains that can be grown, mutated, and assayed, allowing comparative genomic patterns to produce testable hypotheses. Yet no order-wide analysis built on complete genomes of cultured isolates has been available, with the largest published Thermococcales pangenome comprised of ∼30 mostly draft assemblies [15].

Without such analyses, the taxonomy of the Thermococcales has become somewhat disorganised. Genus boundaries within Thermococcales were established when few genomes existed, and 16S rRNA gene identity was used to resolve taxa. 16S rRNA genes are a poor choice for understanding diversity in hyperthermophiles, as the high growth temperature constrains rRNA sequence divergence and shortens branch lengths among closely related lineages [16,17]. Whole-genome metrics such as Mash distance, average nucleotide identity, and average amino acid identity now provide better-calibrated boundaries of both species and genera [18,19], and the Genome Taxonomy Database currently identifies six genera within Thermococcales where classical taxonomy recognizes three [20,21]. Which of these lineages warrant formal recognition, how many species they contain, and whether independent metrics converge on the same taxonomic boundaries remain to be systematically tested.

A second tension within Thermococcales concerns genome architecture. In most prokaryotes, the order of genes on the genome is far from free, being shaped by operon structure, functional linkage, and replication-associated dosage making the chromosomal position of genes itself a selective trait [22–25]. Archaeal chromosomes are also organized spatially by histones, SMC-family proteins, and higher-order domain structure [26,27]. At first glance, Thermococcales appear to buck this trend – their chromosomes seem structurally labile, with inversions and large-scale rearrangements accumulating even between strains of a single species [28–31]. Against that background, however, the positions of essential gene clusters are conserved relative to the origin of replication [24], implying that even ‘scrambled’ chromosomes retain an underlying organization. Whether this extends beyond the core genome to accessory functions that distinguish Thermococcales species from one another has not been examined. Indeed, even the mobile elements thought responsible for genome ‘scrambling’ themselves occupy a small number of chromosomal hotspots due to the site-specific nature of their integrases [29,32,33]. Modern pangenome partitioning approaches present a powerful way to highlight such regions of genome plasticity, identifying modular gene cassettes shared across pangenomic space, rather than being restricted to gene-level presence/absence [34–36].

Here we assemble a dataset of 122 complete genomes of cultured Thermococcales, spanning all publicly available type species, as well as 67 previously unpublished isolates, and analyse them as a single pangenome. We use four independent genome-scale metrics to evaluate genus and species-level taxonomy, resolve conflicting phylogenetic signal by at the base of the order by detecting unstable taxa, and map the distribution of accessory functions and mobile genetic elements. The resulting picture is one of extreme structural plasticity coupled to a strikingly constrained mode of functional diversification – metabolic pathways and mobile elements are gained and lost as intact modules, and they do so at a limited set of genus-specific chromosomal loci.

## Materials and Methods

### Strain isolation/selection

Thermococcales strains were isolated (as described previously [37]) from worldwide hydrothermal vent sites sampled during oceanographic cruises by Ifremer: IRIS [38]; AMISTAD [39]; and BIOSPEEDO [38]. Strains are deposited in the Université de Bretagne Occidentale Culture Collection (UBOCC) (https://www.univ-brest.fr/ubocc/fr).

### Genome sequencing and assembly

Genomic DNA was isolated as described previously [37], and sequenced using a combination of Illumina and Oxford Nanopore Technologies. Paired-end (PE) Illumina libraries were prepared by sonicating DNA to 100-1000 bp using the E220 Covaris Focused-Ultrasonicator (Covaris, Inc.) followed by end-repair, 3’-adenylation, ligation of NEXTflex HT barcodes (Bio Scientific Corporation), and 12 cycles of PCR using the Kapa Hifi Library Amplification kit (Kapa Biosystems). The library was analysed using the Agilent 2100 Bioanalyzer and qPCR quantification (Agilent Technologies), and sequenced on an Illumina MiSeq (2×300 bp). Reads were trimmed using the FastX package to remove low-quality nucleotides (Q<20), sequencing adaptors, and short sequences <30 nt. Preparation of Nanopore libraries differed with available technologies and is outlined in Table S1. Libraries were sequenced with a MiniION device with the MinKNOW basecalling software. Merged Illumina PE reads were assembled with Newbler v2.9 [40]; when Nanopore reads were available, Spades v3.6.0 was used for hybrid assembly [41]. When required, PCR and Sanger sequencing were used to fill gaps. The closed genomes were then deposited in the Microscope mage platform for classical genomes metrics, annotation, and quality checking [42] (Table S2).

### Genome classification and analysis

Genomes were classified taxonomically by combining GTDB-tk [43] with three distance metrics (Fig. S1): i) Mash distance using MinHash (v2.3) dimensionality reduction [44]; ii) global average nucleotide identity (ANI) using FastANI (v1.3) [45]; and iii) average amino acid identity (AAI) using CompareM (v0.1.2) (https://github.com/dparks1134/CompareM). Thresholds for species were set at <0.05 for Mash, >95 for ANI and AAI; and for genera at <0.3 for Mash, >76 for ANI, and >75 for AAI [44,45]. Codon bias analysis was performed in R (v5.5.1) [46] using FactoMineR (v2.11)[47], and biplot visualization was generated with factoextra (v1.0.7)[48] (Fig. 2C).

### Pangenomic and metabolomic analyses

Pangenomes were generated using PPanGGOLiN (v2.0.3) [34] using 60% identity and 60% coverage at the family level, and 70% identity and 60% coverage at the genus level. Representatives from protein families identified with PPanGGOLiN were assigned to KEGG Orthologs using KofamScan [49], and the CLI tool kegg-pathways-completeness [50] used to assign KEGG modules and BRITE categories, as well as pathway completeness. Representative regions of genomic plasticity (determined by PPanGGOLiN) were extracted and aligned with Clinker (v0.0.31)[51] and manually annotated in Adobe Illustrator v25.4.1.

### Phylogenetic reconstruction

Initial phylogenies were generated from a dataset of 76 archaeal core genes [52], and subsets of these comprised ribosomal or non-ribosomal proteins (without the RNA polymerase) using Methanofastidiosales as an outgroup. Phylogenies were generated by the concatenation of the 76 archaeal core proteins with Anvio (v7.1) [53,54], by alignment with MAFFT (v7.490) [55] trimming with trimAI (v1.4.1) [56], and by phylogenetic reconstruction with IQTree (v1.6.12) [57].

Further phylogenies were generated using 142 Thermococcales core proteins selected from the Thermococcales pangenome and also found in at least in 75% of each group’s genomes of the outgroup (12 Hadesarchaea, 12 Methanofastidiosales, and 12 Methanomada), along with respective proteins from 36 outgroup genomes. Each protein family was aligned with MAFFT (v7.490) [55], trimmed with trimAI (v1.4.1)[56], and phylogenies inferred with IQTree (v1.6.12) [57]. MonoPhylo [58] was used to determine the basal genera of each tree. For the final tree, the 142 proteins were concatenated using a partition approach [59] with the best model determined for each individual protein [60]. In both cases the obtained trees were open and annotated with additional information using iTOL.

### Mobile genetic element identification

Episomal plasmids and viruses were identified as extrachromosomal, closed circular assemblies in genome sequencing. Chromosomally integrated elements were identified by integrase-tRNA boundaries, as described previously [32]. Extracted elements were aligned with Clinker (v0.0.31) [51] and manually annotated in Adobe Illustrator v25.4.1

## Results

### Dataset description

Through collaborative sampling, isolation, and sequencing, we generated 67 complete genomes from cultivated members of the archaeal order Thermococcales. Several of the genomes generated in this work have been previously reported in independent studies [29,61–66] while others are described here for the first time. To capture the full described diversity of Thermococcales, our dataset also includes genomes of type strains that had not been sequenced, together with all publicly available Thermococcales genomes available through NCBI and ENA (as of 4th October 2023).

The strains included in this study originate primarily from hydrothermal vent environments spanning multiple ocean basins, including the Pacific, Atlantic, Indian, and Mediterranean Sea (Fig. 1A). A smaller number of isolates were recovered from shallow hydrothermal systems, terrestrial hot springs, and subsurface oil reservoirs, highlighting an ecological breadth that is often underappreciated for Thermococcales (Fig. 1B). Although sampling sites are globally distributed, isolates are disproportionately derived from northern hemisphere vent sites, likely reflecting historical sampling intensity and accessibility rather than true biogeographic restriction. Indeed, 21 isolates in our dataset originate from southern hemisphere sites, including 2 freshwater Thermococcales isolated from terrestrial geothermal springs in New Zealand [67–69]. Key genomic features of all strains are summarized in Table S3. Genome sizes range from 1.52 to 2.38 Mbp, congruent with the compact genomes characteristic of hyperthermophilic archaea [70]. All genomes encode between 45 and 49 tRNA genes and contain a single rRNA operon, with the exception of *Palaeococcus pacificus* DY20341, which harbors two distinct 16S rRNA loci. The total number of predicted protein-coding genes varies from 1599 to 2628 per genome. Despite their phylogenetic relatedness and shared hyperthermophilic lifestyle, Thermococcales exhibit substantial variation in nucleotide composition, with G+C content ranging from 39.4% to 56.5%. No clear relationship was observed between overall high G+C content, genome size, or geographic location, reinforcing previous observations that elevated G+C content is not a requirement for DNA stability in hyperthermophilic organisms [71]. We could observe that the *Pyrococcus* strains, known to have a higher optimal temperature than *Thermococcus* strains, harbored a lower G+C than these *Thermococcus* strains (Table S2, also in Fig. S2).

**Fig. 1:**
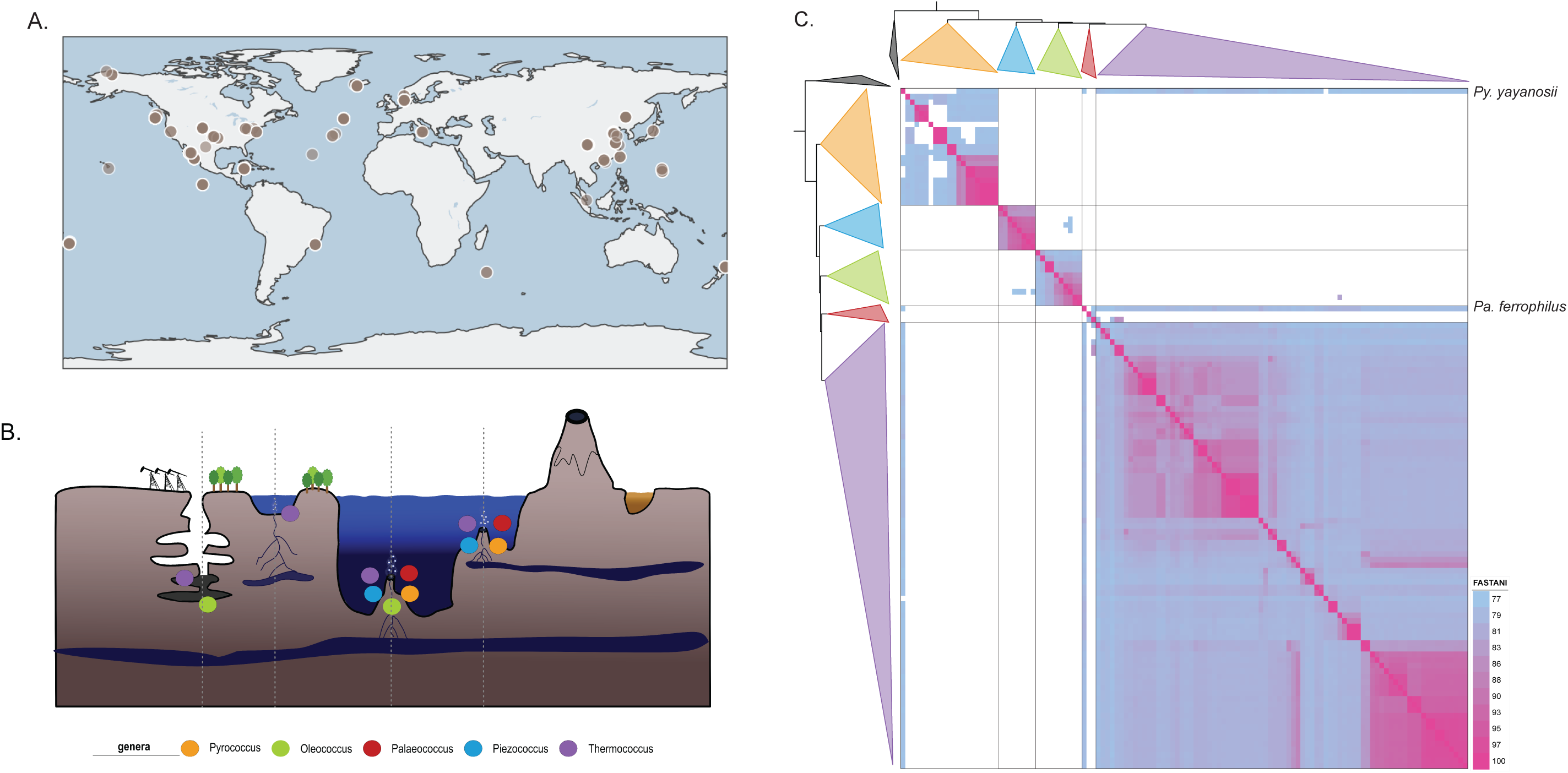
Diversity of Thermococcales strains across geography and habitats. A-Global map of sampling sites. The location of Thermococcales signal on metagenomic data or the location of strain genomes available on the sites of the NCBI is indicated by dots. **B-The diversity of habitats of our complete set of Thermoccales genomes.** In each habitat presence of each genus is indicated by a colored dot. The orange corresponds to Pyrococcus, the green to Oleococcus, the red to Palaeococcus, the blue to Piezococcus and the purple to Thermococcus. **C-Heatmap visualization of the diversity of Thermoccales genus** using whole genomes ANI values. For Average nucleotide identity (ANI) analysis, only values greater than 77 are shown; the higher the value, the closer the genomes are. If the ANI between two genomes is above 76, they belong to the same genus; if it’s above 95, they belong to the same species [45]. (Heatmap of ANI, AAI and Mash are provided as Sup Fig. 1)

### Comparative genomics supports a revised genus-level classification of Thermococcales

In the current release of GTDB (10-RS226) [20,21], Thermococcales, together with Methanofastidiosa [72], are placed within the phylum Methanobacteriota B (confusingly, Methanobacteria are not part of this phylum, instead occupying Methanobacteria A). In parallel with recent efforts to stabilize archaeal nomenclature, we favor adoption of the placeholder phylum name Acherontiarchaeota, reflecting the original higher-level lineage Acherontia proposed previously [8].

At the genus level, Thermococcales taxonomy has historically recognized three genera: *Pyrococcus*, *Palaeococcus*, and *Thermococcus*. However, comparative studies have suggested that this classification underrepresents the true genomic diversity of the order [73], a conclusion echoed by GTDB which currently delineates six genus-level lineages (Pyrococcus, Palaeococcus, Thermococcus, Thermococcus_A, Thermococcus_B, and Thermococcus_C represented by a single MAG of *Thermococcus sp.* JdFR-02). To systematically evaluate genus- and species-level structure within our expanded Thermococcales dataset, we applied a suite of complementary genome-scale metrics.

Although 16S rRNA gene sequence identity remains widely used for prokaryotic taxonomic delineation [74,75], this marker is poorly suited for Thermococcales. Hyperthermophilic growth constraints are known to limit 16S rRNA sequence divergence, resulting in compressed branch lengths and reduced resolving power among closely related lineages [16,17]. In addition, Thermococcales genomes frequently exhibit large-scale rearrangements and nested inversions, likely driven by mobile genetic elements [24,28,29], complicating approaches that rely on conserved genome architecture for taxonomic inference such as hybridization.

We therefore adopted a multi-metric genome-wide strategy combining: 1) Mash distance – a MinHash-based estimator of genetic divergence [44]; 2) whole-genome average nucleotide identity (ANI) computed with FastANI [45]; 3) average amino acid identity (AAI) across shared orthologues; 4) taxonomic assignation using GTDB-tk [43,76]; and 5) phylogenetic placement based on concatenated conserved protein markers [52]. Across all four approaches, Thermococcales genomes segregated into five well-supported genus-level clusters (Fig. 1C). These five lineages correspond to the genera in GTDB named *Pyrococcus*, *Palaeococcus*, *Thermococcus*, *Thermococcus A*, and *Thermococcus B*. To reflect ecological and physiological distinctions within these latter two groups, we propose renaming *Thermococcus A* as “*Oleococcus*” referencing the isolation of several members currently classified as *Thermococcus* from subsurface oil reservoirs e.g. *T. sibiricus* [77]*, T. bergensis* [78] and the genomically incomplete *T. petroboostus* 101 C5 [79,80]; and *Thermococcus B* as “*Piezococcus*”, reflecting the piezophilic nature of this lineage, exemplified by *T. barophilus* [81]. As multiple genera share the same initial letter, two-letter abbreviations are used throughout the manuscript when referring to individual species (*Py. = Pyrococcus, Pa. =Palaeococcus, Pi. = Piezococcus*).

Notably, *T. atlanticus* occupies a deep branch comparable to the genus-level divisions defined above. However, as this lineage is represented by a single genome, we refrain from proposing additional genera until broader sampling of related strains allows for confident taxonomic resolution.

This revised genus-level classification resolves the paradoxical placement of *Palaeococcus* within the broader *Thermococcus* clade under previous taxonomies and corrects the conflation of multiple deeply branching lineages under a single genus name.

### Genome-wide metrics delineate species boundaries

Species-level delineation within Thermococcales proved more variable than genus-level classification, with modest differences observed between genome-based metrics (70 species defined by 95% ANI, 68 by 95% AAI, and 66 by <0.05 Mash distance) (Fig S1, Table S3). These discrepancies largely reflect borderline cases where two of the three metrics agree close to borderline thresholds rather than fundamental disagreements among methods.

Sampling across genera was uneven, with the genus *Thermococcus* substantially overrepresented relative to other lineages. In contrast, *Palaeococcus* is represented by only two complete genomes (*Pa. pacificus* [82] and *Pa. helgesonii* [83]), and a single representative partial genome (*Pa. ferrophilus* [84]). Within *Pyrococcus,* genome-based analyses support 10 distinct species-level clusters. Seven correspond to previously described species (*Py. abyssi* [85], *Py. endeavori, Py. furiosus* [86]*, Py. horikoshii* [87]*, Py. kukulkanii* [65]*, Py. yayanosii* [88,89], and *Pyrococcus chitonophagus* (previously described as *Thermococcus chitonophagus* [90]), while three represent new species (*Py. sp.* EXT15c, *Py. sp.* IRI42c, *Py. sp.* NA2).

The newly proposed genus *Oleococcus* comprises seven species, encompassing strains previously assigned to *Thermococcus,* including *T. aggregans, T. aegaeus, T. argininiproducens, T. litoralis, T. sibiricus, T. bergensis,* and *T. sp. 2319x1.* All genome-scale comparisons consistently support their separation from *Thermococcus* (Fig. S1) and these species should thus be renamed *Oleococcus aggregans,* etc.

The proposed genus *Piezococcus* comprises 5-6 species, depending on the metric used. These include the canonical *T. barophilus* containing type strain MP, as well as strains CH1 and CH5, and distinct lineages corresponding to *T. paralvinellae, T. sp.* SY098, and *T. barophilus* DT4 and 423_3_23 which require re-naming to differentiate these from true *T. barophilus* strains. One additional strain (*T. sp.* CIR10a) occupies a borderline position and remains unresolved at the species level, grouping with *T. barophilus* 423_3_23 in both Mash and AAI analyses but remaining distinct with ANI (Table S3). As above, these species should be renamed *Piezococcus barophilus,* etc.

The genus *Thermococcus* is the most species-rich genus in the dataset, comprising 42 species. Of these, a subset corresponds to previously described species, including *T. atlanticus, T. barossi, T. camini, T. celer* [91]*, T. celericrescens* [92]*, T. cleftensis, T. coalescens, T. eurythermalis, T. fumicolans, T. kodakarensis, T. nautili, T. onnurineus, T. pacificus, T. peptonophilus, T. piezophilus, T. profundus, T. radiotolerans* [93]*, T. siculi, T. stetteri, T. thioreducens, T. thermotolerans,* and *T. zilligii.* In addition, our analyses support the definition of 16 previously unrecognized species within *Thermococcus* (Table S3). Conversely, several historically proposed species require revision to the level of strains.

Notably, the type culture of *T. prieurii* was found to represent a stable co-culture comprising two strains that resolve as different species across all genome-based metrics. Only one strain harbors the virus TPV1 and plasmids pTP1 and pTP2 [94,95], whereas the other is virus and plasmid-free. The published 16S rRNA gene sequence associated with *T. prieurii* (Genbank FN396976) derives from the virus-free lineage. To avoid continued taxonomic ambiguity, we propose renaming the virus-free strain *T. komboloii,* and reassigning the virus-containing strain as *T. henrietti* strain TPV, consistent with its placement by Mash and AAI metrics. Additionally, we identify several cases of synonymy among described species: strains *T. eurythermalis* A501 [96] and *T. marinus* EJ1 resolve as a single species; as *T. marinus* was described first [93], *T. eurythermalis* should be reassigned to *T. marinius* strain A501. Similarly, *T. celericrescens* TS2 (DSM 17994) and *T. aciditolerans* SY113 form a single species across all analyses, supporting the renaming of *T. aciditolerans* to *T. celericrescens* strain SY113.

A summary of all proposed genus and species names is available in Table S3. Together, these results resolve a complex and uneven species landscape within Thermococcales, establishing a clear species framework on which subsequent evolutionary analyses are built.

### Core genome-derived datasets resolve conflicting phylogenies

To reconstruct evolutionary relationships within Thermococcales, we inferred phylogenies using three marker protein datasets: 1) ribosomal proteins; 2) non-ribosomal proteins; 3) a standard archaeal marker set comprising 76 conserved proteins [52]. While all 3 approaches recovered the genus-level groupings described above, they differed in their resolution of inter-genus relationships, particularly with respect to the basal position of Thermococcales lineages (Fig. S3-6).

Across these initial analyses, either *Pyrococcus* or *Palaeococcus* was recovered as the earliest-branching genus, depending on the marker set used, indicating conflicting phylogenetic signal at the base of the Thermococcales tree (Fig. S3-6). To increase the resolution and reduce stochastic effects associated with small marker sets, we expanded the analysis to genome-wide conserved proteins identified through pangenomic analysis. Core gene families identified through pangenomic analysis were used here solely as a source of conserved phylogenetic markers; broader pangenome structure and functional dynamics are analyzed separately below. From 483 Thermococcales core gene families, we selected 142 proteins also present in at least 75% of outgroup taxa (Methanofastidiosales and Methanomada). Single protein phylogenies were evaluated using MonoPhylo to assess the stability of inferred clades and to identify taxa with inconsistent phylogenetic placement. This stability analysis identified 3 taxa: *Py. yayanosii, Pa. ferrophilus,* and *T. atlanticus* as phylogenetically unstable, exhibiting variable placement across single-gene trees and disproportionately reducing topological support (Fig S7). Exclusion of these rogue taxa increased the frequency with which the proposed genera (*Oleococcus, Piezococcus, Pyrococcus, Palaeococcus,* and *Thermococcus*) were recovered as monophyletic groups across single gene trees (Fig 2A).

**Fig. 2:**
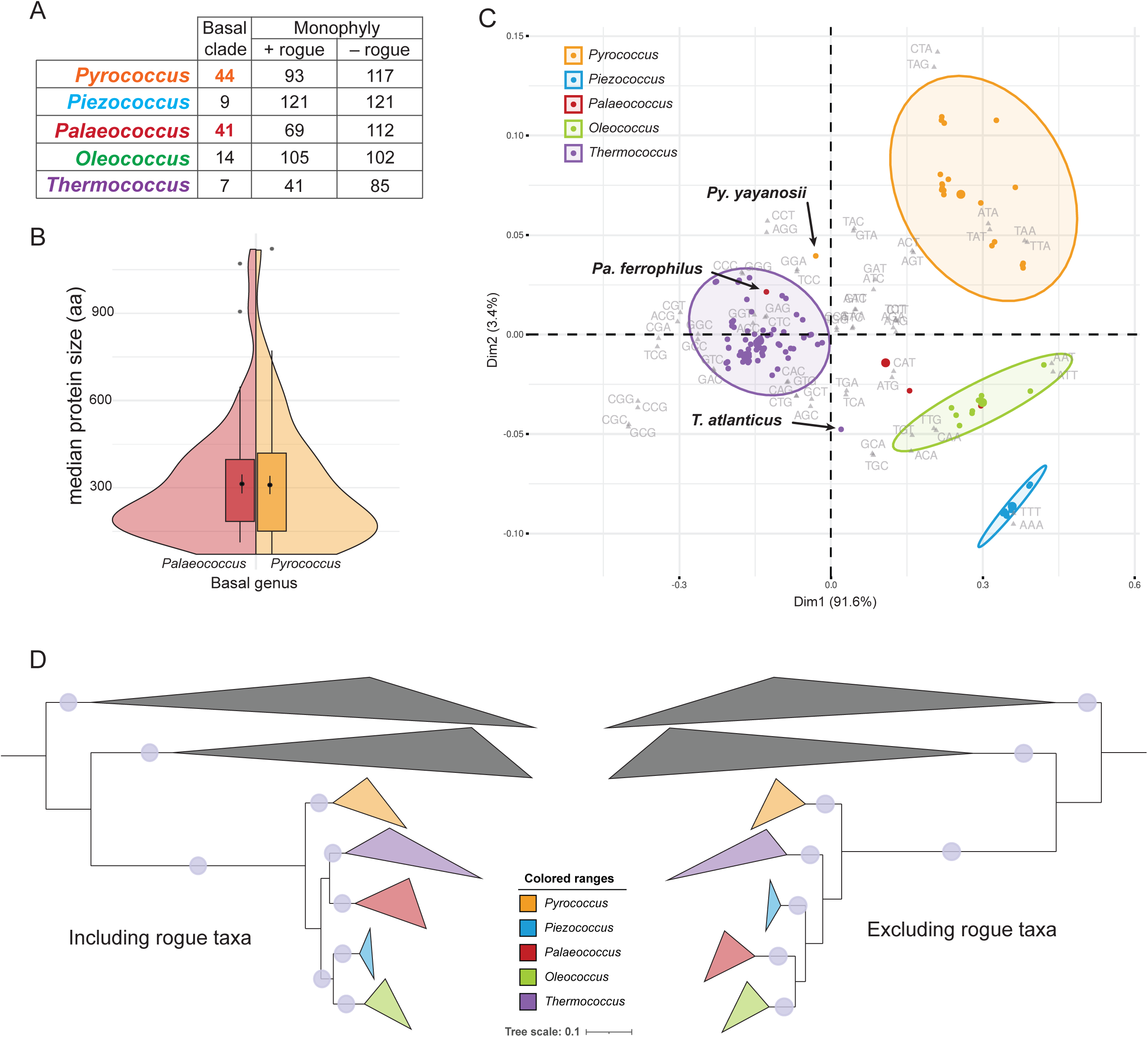
Influence of Rogue Taxa on Phylogenetic Reconstruction and Basal Signal. **A-Summary table** showing, for each single-marker phylogeny, the number of trees in which the investigated genera were recovered as monophyletic and positioned at the base of the phylogeny. The impact of rogue taxa on genus monophyly is also indicated. **B-Split-violin plot** comparing the size distribution of proteins whose phylogenetic signal supports either *Palaeococcus* or *Pyrococcus* as the basal genus. C-**D-Collapsed phylogenic tree of the Thermococcales build with 142 core proteins with or without the rogue taxa** - on an alignment of 39788 positions by using a partition approach with the best model determined before for each individual proteins. For both phylogenies, nodes supported by an ultrafast bootstrap value >95 and an SH-aLRT support value >80 are marked with light blue dots.

The unstable behavior of these genomes may reflect extensive horizontal gene transfer of core genes with other Thermococcales lineages or, alternatively, their status as sole representatives of deeply branching, unrecognized genera. Coherent with the former hypothesis, ANI analysis revealed evidence of gene sharing between *Py. yayanosii* and *Thermococcus*, and between *Pa. ferrophilus* and other lineages (Fig S1). In addition, codon usage analyses resolve clear genus-specific clusters, yet place these rogue genomes intermediate between their assigned genera and *Thermococcus* (Fig 2C). Their genomic GC contents are also discordant with the surrounding species in their respective genera (Fig S2), reinforcing that these species have experienced distinct evolutionary trajectories relative to their closest classified relatives.

Examination of single-gene trees confirmed that support for either *Pyrococcus* or *Palaeococcus* as the basal Thermococcales lineage was distributed across many loci, with 44 proteins supporting *Pyrococcus* and 41 supporting *Palaeococcus* as basal (Fig 2A). Separating marker genes into ribosomal and non-ribosomal sets did not preferentially support one basal topology over the other (for ribosomal 12 and 17 and for non-ribosomal 32 and 24, for pyrococcus and palaeococcus respectively). Likewise, markers supporting *Pyrococcus* or *Palaeococcus* as the basal lineage were not skewed by length (Fig 2B). As longer proteins typically carry stronger phylogenetic signal, we might expect longer proteins to more accurately resolve evolutionary histories. The absence of such patterns suggests that the alternative basal placements reflect deep evolutionary signals rather than methodological artefacts. To maximize phylogenetic resolution, we next constructed phylogenies using a concatenation of the full 142-protein dataset, both including and excluding the identified rogue taxa. With this increased signal, *Pyrococcus* was recovered as the basal genus, and the monophyly of the five genera was strongly supported regardless of the presence of rogue taxa (Fig 2D). Notably, inclusion of rogue taxa did affect the placement of *Palaeococcus,* while the relative relationships among all other taxa remained unchanged. This sensitivity likely reflects sparse sampling within *Palaeococcus* (only three genomes, including one rogue species), rendering its placement less stable. Resolving the evolutionary position of this lineage will require additional genomes, as well as improved sampling of deeply branching taxa such as *T. atlanticus,* which may ultimately warrant recognition as a distinct genus. A full species tree inferred from the concatenated 142-protein dataset is available in Fig S8.

### Thermococcales pangenome reveals extensive genomic plasticity

Comparative genomic analysis of the complete Thermococcales dataset using PPanGGOLiN reconstructed a pangenome comprising 264,322 genes organized into 21,490 gene families (Fig. 3A). Partitioning of these families revealed a relatively small persistent genome (1,311 families, 6.1%), alongside a larger shell (2,513 families, 11.7%) and a substantial cloud component (17,666 families, 82.2%) (Supp. Dataset. 1 – all gene families separated by persistent-shell-cloud), indicating extensive variability in gene content across the order. The persistent gene families predominantly encode information processing pathways, including ribosomal proteins, tRNA synthetases, RNA polymerase subunits, and DNA replication and repair machinery, as well as central metabolic pathways such as glycolysis and archaeal lipid synthesis [76].

**Fig. 3:**
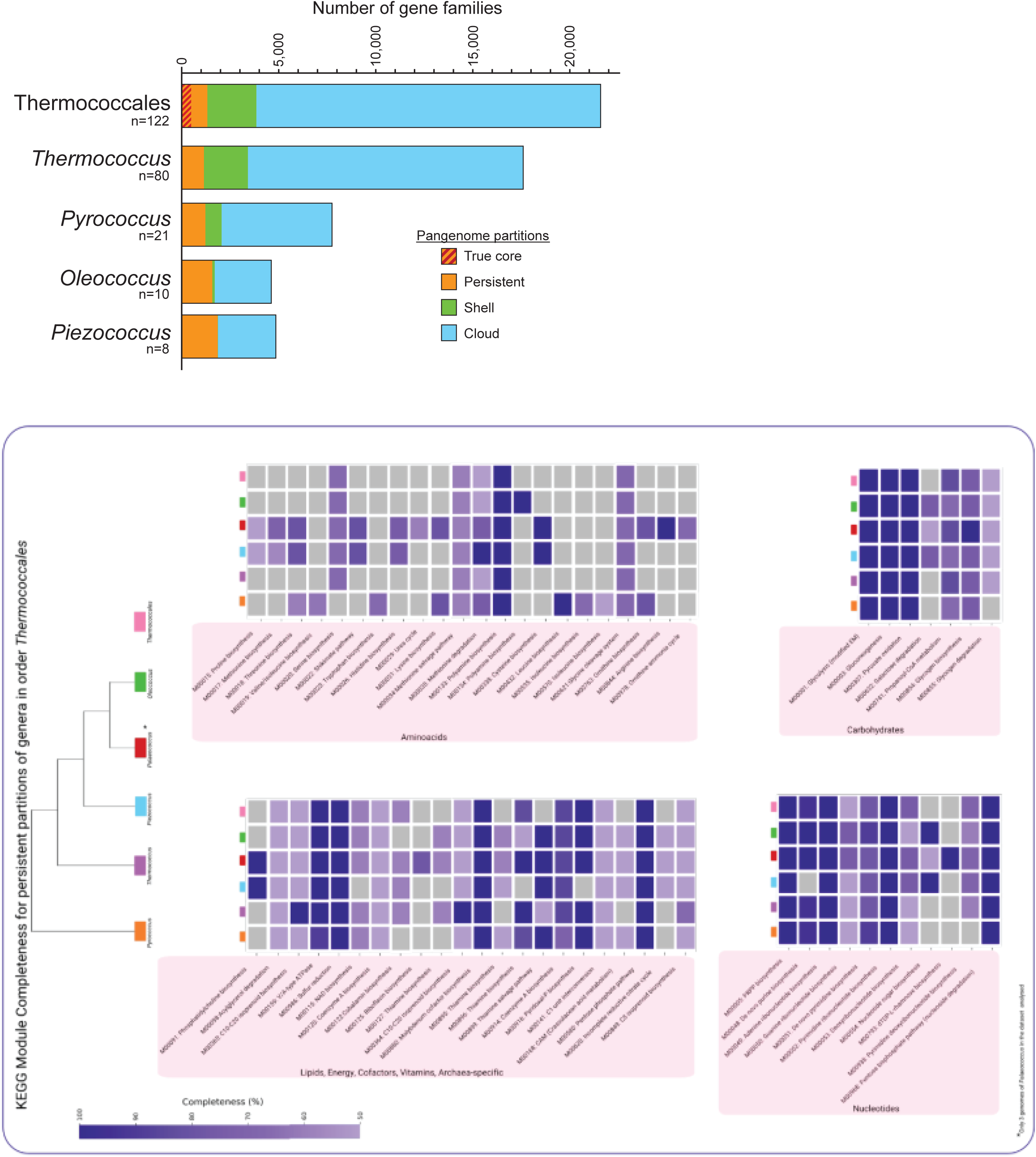
Comparative pangenomic analysis of Thermococcales and individual genera. **A-**Bar plots showing the number of gene families in each pangenome partition for the entire *Thermococcales* order and for the genera *Pyrococcus*, *Thermococcus*, *Oleothermococcus*, and *Piezococcus*. Persistent, shell, and cloud gene families are shown in orange, green, and blue, respectively. **B-**Heatmap illustrating the completeness of predicted metabolic pathways within the persistent genome fraction of *Thermococcales* and the different genera. Values correspond to the proportion of pathway-associated genes detected in each persistent genome.

Among the shell and cloud families, 2,161 gene families are organised into 496 conserved modules, defined as sets of genes that are colocalized across genomes. These conserved modules likely correspond to functional units within the pangenome, including pathways involved in metabolism, stress response, and mobile genetic elements (MGEs), and highlight that a significant fraction of the Thermococcales genome variability is structured rather than random.

At the level of individual genomes, the strict core genome represents a median of 23% of encoded genes (483 gene families). When considering the broader persistent partition, a typical Thermococcales genome is composed of approximately 55% persistent, 27% shell, and 17% cloud genes (Fig. S2). Although only 6.1% of gene families across the order belong to the persistent partition, these families account for over half of the genes within any individual genome. In contrast, cloud gene families comprise 82.2% of all gene families identified, yet contribute only 17% of genes per genome. This asymmetry illustrates that each genome is dominated by a large shared backbone of conserved functions, while the variable portion of each genome draws from a much larger accessory gene pool, providing substantial capacity for metabolic and ecological differentiation. Consistent with this observation, the Thermococcales pangenome is open, such that the addition of new genomes is expected to continue expanding the total gene repertoire. This is further illustrated by genus-level pangenomes, where expanded sampling from 21 *Pyrococcus* to 80 *Thermococcus* genomes, and ultimately to all 122 Thermococcales genomes, results in minimal change in the size of the persistent gene set, whereas the number of cloud gene families increases substantially (Fig. 3A). Thus, additional genomes primarily contribute novel accessory gene families rather than expanding the conserved core. Notably, our pangenome mirrors earlier analyses based on ∼30 genomes [15] demonstrating that even with a 4-fold increase in sampling, gene diversity continues to accumulate rather than plateau.

### Archaella and Type IV pili are conserved

We noted that genes associated with the archaellum were found in the persistent gene set. To assess the conservation of these systems across Thermococcales, we identified all archaellum structural genes (*flaB-J*) and mapped their presence onto the phylogenetic tree (Fig. 4). Among the ∼70 defined Thermococcales species, only 13 lack evidence of an archaellum. Notably, of the two *T. zilligii* strains, only one encodes a complete archaellum operon, while the strain previously designated *T. waiotapuensis* is missing 7 of the 8 required genes. The near-ubiquitous distribution of archaella across Thermococcales underscores their importance in motility and surface attachment [97], and also supports their presence in the last common ancestor of the order.

**Fig. 4:**
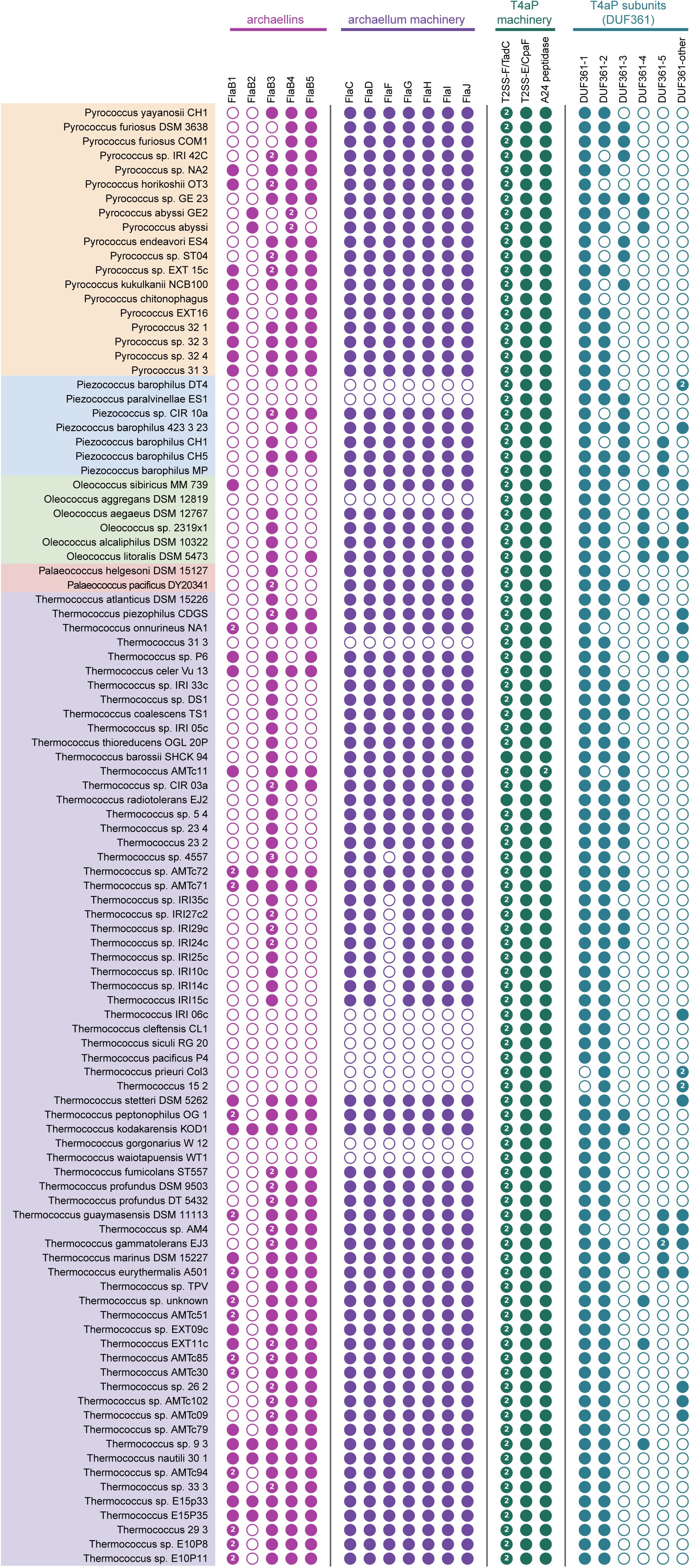
Identification and Distribution of archaellum and Type IVa pili (T4aP) components across Thermococcales genomes. Colored dots indicate the presence (filled) or absence (empty) of each component. Genomes are grouped by genus and displayed using the color scheme defined above: *Thermococcus* (purple), *Pyrococcus* (orange), *Oleothermococcus* (green), and *Piezococcus* (blue).

Interestingly, the archaellum genes are organized in a single operon but encode varying numbers of FlaB paralogs, ranging from a single copy in *Pa. helgesoni* to six copies in *T. sp.* AMTc71 (Fig. 4). FlaB constitutes the primary structural component of the archaellum, forming the filament [79]. In *T. kodakarensis,* all five paralogs have been detected in purified archaella, albeit in different proportions [80], suggesting that multiple archaellins can co-assemble into a single filament. In haloarchaea, two archaellins (FlaA and FlaB) similarly contribute to the filament structure, with their relative abundance varying under different growth conditions, particularly salinity, consistent with functional diversification of paralogs [81, 82]. *Methanococcus maripaludis* encodes three *flaB* genes, and deletion of any individual paralog results in defects in archaellum assembly or motility, indicating these proteins have non-redundant contributions [83]. Despite these observations, the functional significance of maintaining multiple FlaB paralogs in most Thermococcales species remains unclear, particularly given that a single *flaB* gene is sufficient in several species.

The original description of *T. waiotapuensis* reported the presence of “flagellated cells” by electron microscopy [60]. However, we did not detect a complete archaellum operon in our pangenome analysis of this strain (Fig 4). This discrepancy raises the possibility that at least some filaments observed in Thermococcales do not correspond to canonical archaella. In archaea, additional type IV pili systems have been well characterized in *M. maripaludis* [98] and Crenarchaea [99], and are also predicted in Thermococcales species (including *T. waiotapuensis*) based on the presence of multiple prepilin genes and associated processing enzymes such as EppA [86]. We therefore examined Thermococcales genomes for genes encoding type IV pili systems. Interestingly, pili systems were found in every genome (Fig. 4), suggesting they may be even more important to Thermococcales physiology than archaella, and were likely present in the last common ancestor of Thermococcales. This is reinforced by the concordance between prepilin peptidase paralogs and the five Thermococcales genera, with further divergence seen between *Thermococcus* clades.

### Metabolic diversity arises from gain and loss of biosynthetic modules

Across Thermococcales, pathways underlying central carbon metabolism and energy conservation are conserved, forming part of the persistent genome. All examined genomes encode enzymes of the archaeal-specific variants of glycolysis (modified Embden-Meyerhof pathway) and gluconeogenesis, including ADP-dependent kinases [6] and glyceraldehyde-3-phosphate:ferredoxin oxidoreductase [7], together with pyruvate oxidation, the pentose bisphosphate (PBP) pathway [100] and sulfur reduction mechanisms (Fig. 3B). Core nucleotide biosynthesis pathways, including adenosine and pyrimidine ribonucleotide synthesis, as well as molybdenum cofactor biosynthesis, are likewise retained across all genera. In addition, all examined genomes encode complete polyamine biosynthesis pathways [101] (Fig 3B), consistent with the proposed role of stabilizing nucleic acids [102,103] and proteins [104] under extreme thermal conditions [105]. Collectively, the conservation of these core pathways across genera indicates strong purifying selection on metabolic functions essential for growth in extreme conditions.

In contrast to this conserved metabolic backbone, Thermococcales display substantial diversity in nutrient utilization and biosynthetic capabilities. This asymmetry is particularly evident in carbohydrate utilisation and amino acid requirements observed in culture. For example, growth assays across multiple Thermococcales species have revealed differences in the ability to utilise both simple and complex carbohydrates that are decoupled from cellular phylogeny [106]. Similarly, comparative physiological studies indicate that amino acid requirements vary extensively between species, with some exhibiting strict auxotrophies while others retain biosynthetic capabilities [67,85,107].

Genome-scale analysis reveals that these phenotypic differences likely correspond to the presence or absence of complete biosynthetic modules rather than gradual pathway erosion. Prediction of regions of genome plasticity (RGPs) across Thermococcales genomes shows that pathways for histidine, lysine, aromatic amino acids (shikimate pathway), and tryptophan biosynthesis, together with peptide and amino acid transport systems, are organised into discrete gene clusters that are either fully present or entirely absent from individual genomes (Fig. 5). The presence of these modules provides a strong predictor of biosynthetic potential at the genomic level; however, translating pathway presence or absence into observed prototrophy or auxotrophy is not straightforward. *Pi. barophilus*, for example, requires only 3 amino acids for growth at low pressure but becomes auxotrophic for 16 under high hydrostatic pressure [108] indicating that pathway activity can be condition-dependent despite completeness at the gene level.

**Fig. 5:**
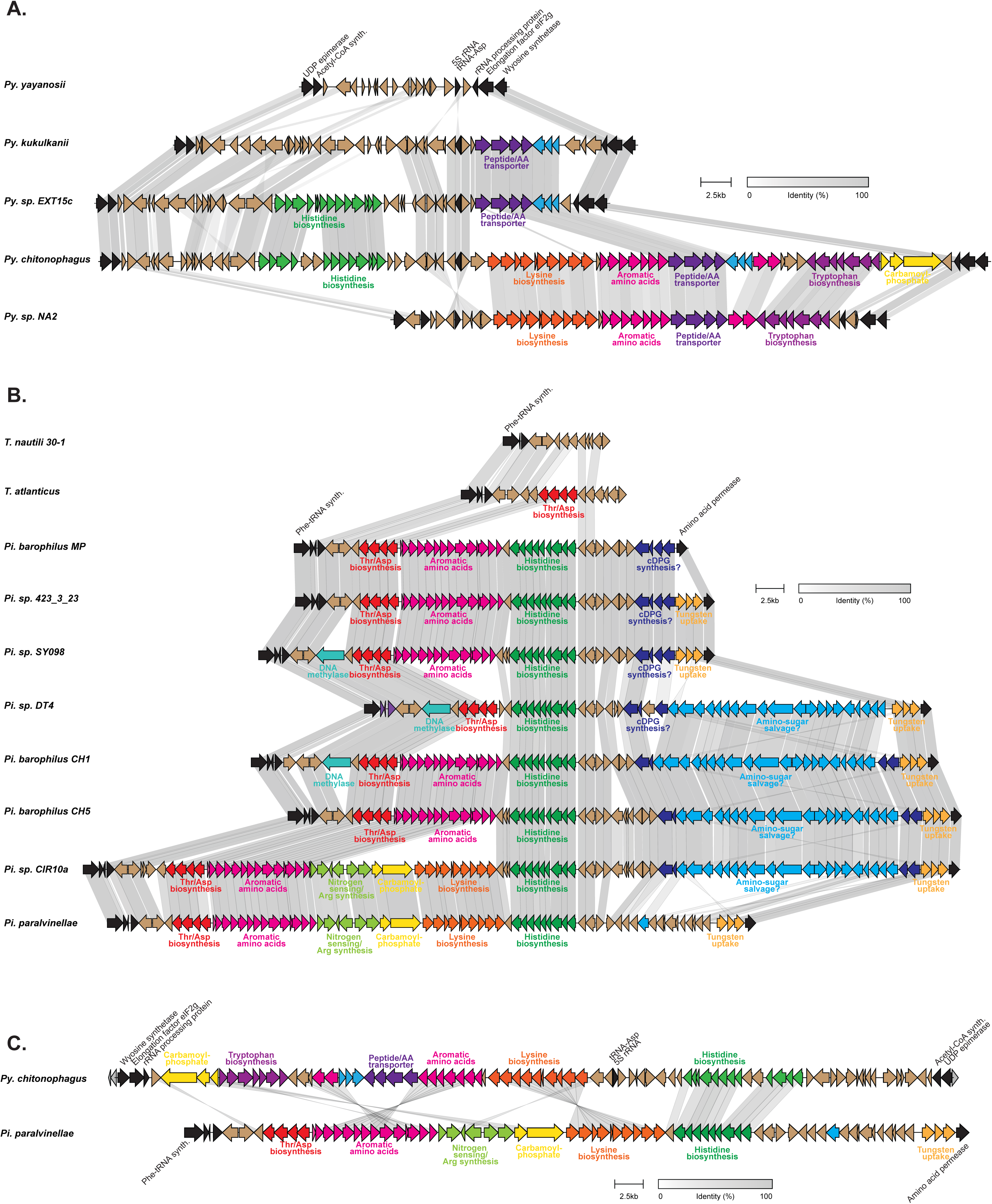
Synteny and functional diversification of metabolic genomic regions in Pyrococcus and Piezococcus. Comparative synteny analysis of a variable genomic region among selected *Pyrococcus* genomes (A) and in *Piezococcus* and Thermococcus genomes (B). Arrows represent predicted coding sequences, with colors indicating genes associated with specific metabolic functions. Homologous genes are connected and colored according to their amino acid identity. The region contains genes involved in histidine, lysine, aromatic amino acid, and tryptophan biosynthesis, as well as peptide/amino acid transport and carbamoyl phosphate metabolism. Scale bars correspond to 2.5 kb.

Notably, although homologous biosynthetic modules are found across *Pyrococcus* and *Piezococcus* species, their chromosomal integration sites differ between genera. In *Pyrococcus,* variable biosynthetic and transport pathways are embedded within a conserved locus flanking the 5S rRNA and tRNA^Asp^ genes, bounded by UDP epimerase and wyosine synthetase (Fig 5A). In contrast, closely related modules in *Piezococcus* species are located at a distinct chromosomal region centered on the *folP* to *pheT* locus, often extending upstream of *folP* (Fig 5B). Direct comparison of representative loci from *Pyrococcus* and *Piezococcus* demonstrates that while the surrounding chromosome context differs between genera, the gene content and internal organization of these modules are conserved (Fig 5C). This conserved genetic positioning, coupled with all-or-nothing pathway presence, is consistent with metabolic diversity arising through gain and loss of intact functional modules rather than piecemeal gene turnover. Such modularity likely results from frequent acquisition of multi-gene loci, indicating that horizontal transfer has played a central role in shaping the metabolic repertoires of Thermococcales. Indeed, phylogenetic reconstruction of the general nucleoside ABC transporter (K23537, K23535, K23536 [module 63]) reveals this module likely has different evolutionary histories in *Pyrococcus* compared to the other Thermococcales genera, being most closely related to that of Crenarchaeota from hydrothermal vent-associated Crenarchaeota such as *Staphylothermus marinus* and *Candidatus* Tiamatella incendiivivens [109,110], than with those from other Thermococcales, suggesting acquisition by horizontal gene transfer (Fig S9).

### Diverse Thermococcales MGEs differentially distribute across lineages

In addition to the 122 Thermococcales chromosomes, we identified a diverse mobilome comprising 42 extrachromosomal elements and 72 integrated MGEs, including plasmids and viruses (Fig. 6). These elements range from small cryptic plasmids to large replicons and integrated viruses, underscoring the extensive contribution of MGEs to Thermococcales genome evolution.

**Fig. 6:**
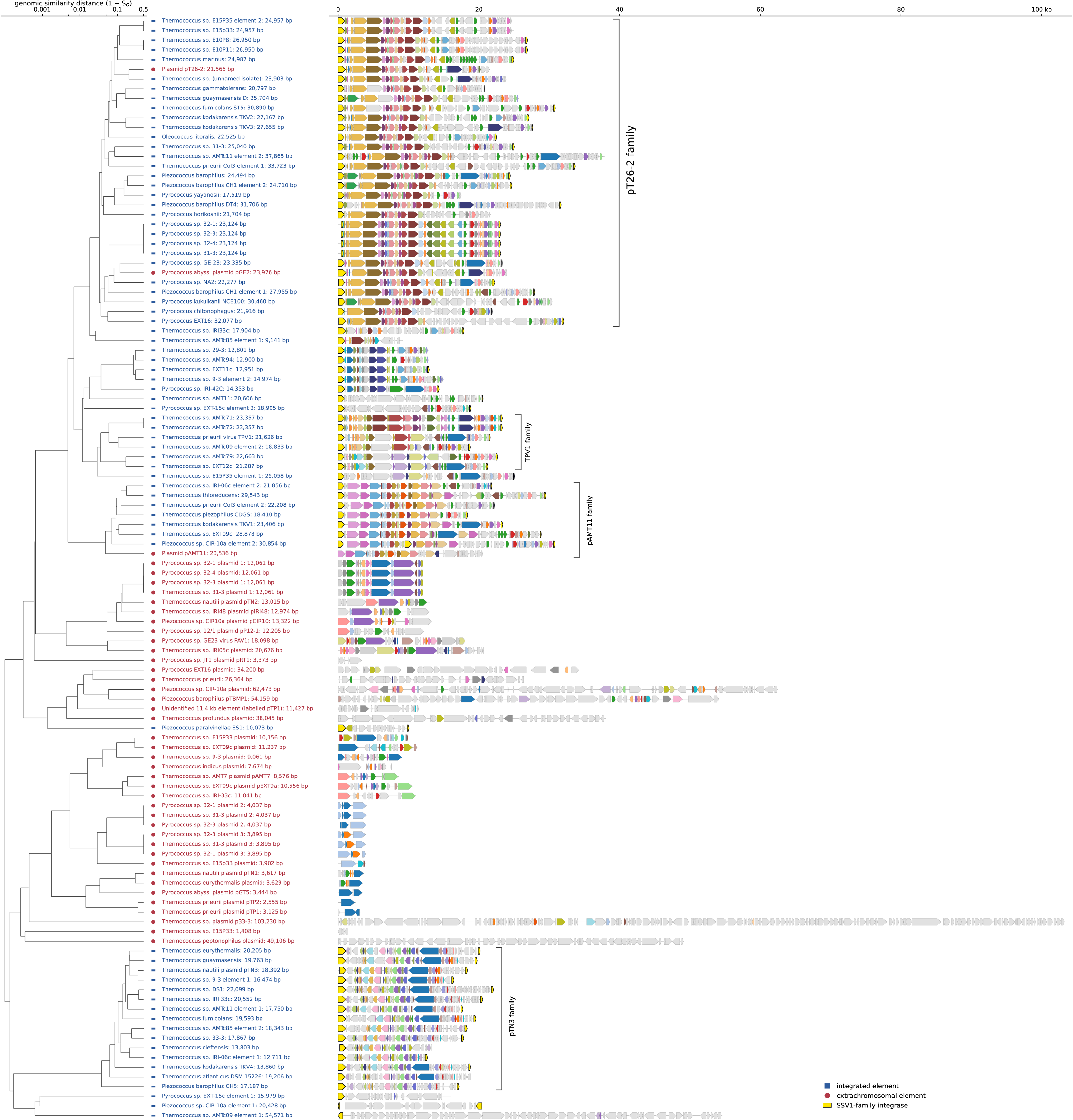
Diversity and distribution of extrachromosomal and integrated mobile genetic elements in the Thermococcales. Elements were clustered according to genomic similarity (1 − Sørensen distance). The dendrogram shows relationships among integrated (blue dots) and extrachromosomal (red dots) elements. Gene maps display the organization of homologous protein families, with SSV1-family integrases highlighted in yellow. Major element families (pT26-2, TPV1, pAMT1, and pTN3) are indicated.

Small cryptic plasmids (∼1.4–4.0 kbp) constitute a prominent fraction of the extrachromosomal elements identified. Such plasmids have historically been instrumental in the development of shuttle vectors for Thermococcales genetics, including pTN1 from *T. nautili* [111], pTP2 from *T. prieuri* [112], and pGT5 from *Py. abyssi* [113], which together enable heterologous transformation of and protein expression in multiple species. Our dataset expands this repertoire substantially. Several newly identified plasmids with little or no detectable similarity to known replication modules (Fig. 6), including two coexisting plasmids in *Py. sp.* 32-1 (3.9 and 4.0 kbp), the smallest plasmid in the dataset from *T. sp.* E15P33 (1.4 kbp), and pRT1 from *T. sp.* JT1 [114]. The novelty of these replicons suggests untapped potential for expanding the genetic toolkit of Thermococcales, particularly in species that are readily manipulated at the chromosome level but for which stable shuttle vectors have remained elusive, such as *T. onnurineus* [4]*, Pi. barophilus* [1], and *Py. yayanosii* [5].

Plasmids of intermediate size (∼7–13 kbp) have previously been described in several Thermococcales species, including pTN2 and its relatives, as well as pCIR10, pIRI48, and pAMT7 [115,116]. Our dataset reveals additional diversity within this size class, including a new member of the pTN2 family (pP32-1 from *Py. sp.* 32-1) and several other intermediate-length plasmids with limited homology to known elements (Fig. 6). These findings suggest that intermediate-sized plasmids represent an active evolutionary reservoir within Thermococcales, maintaining diverse replication and maintenance strategies without obvious specialization for conjugation of virion production. Although their biological roles remain unclear, the consistent recovery of intermediate-sized plasmids across multiple genomes indicates that this class of replicons is a stable and recurring component of Thermococcales mobilomes.

A smaller number of large plasmids have previously been identified, including the 54 kbp pTBMP1 from *Pi. barophilus* MP [117], a 38 kbp plasmid from *T. profundus,* and the characterized conjugative plasmid pT33-3 (103 kbp) from *T. nautili* strain 33-3 [62]. Our dataset adds two additional large plasmids, pCIR10a (62 kbp) and pEXT16 (34 kbp), which share few genes with known Thermococcales MGEs. While pT33-3 is capable of conjugative transfer, other plasmids lack clear hallmarks of infectivity and encode a mixture of metabolic and information-processing genes.

Even more prevalent than extrachromosomal elements are integrated MGEs embedded within Thermococcales chromosomes. By identifying conserved N- and C-terminal fragments of SSV1-like integrases [29,32], we detected 72 integrated elements across the dataset (Fig. 6). Most could be classified based on similarity to previously described Thermococcales MGEs, including the plasmids families pT26-2 [118], pAMT11 [119], and the integrative virus-like elements TPV1 [94] and pTN3 [91]. Elements of the pT26-2 family are particularly abundant [118], infecting members of all Thermococcales genera except *Palaeococcus*, and frequently occur in multiple non-identical copies within a single genome. Notably, the ‘virus-like’ regions of the *T. kodakarensis* genome, TKV2 and TKV3, correspond to integrated pT26-2 family plasmids [120]. In contrast, pAMT11 family elements are less common (n=8), and include the TKV1 in *T. kodakarensis* [120].

Integrated virus families show sharply different distributions within Thermococcales. TPV1-like elements concentrate within a single *Thermococcus* subclade, including *T. prieurii, T. nautili,* and *T. sp.* AMTc30, while pTN3-like elements are distributed broadly across *Thermococcus* yet remain absent from the other genera. Synteny analysis further reveals a close relative of PAV1 in *T. sp.* IRI05c. Virion production has so far been reported only for PAV1 in *Py. sp.* GE23 [121] and TPV1 in *T. prieurii* [94], but the presence of an additional PAV1-like element and 15 integrated pTN3-like elements encoding putative capsid proteins [122] raises the possibility that additional Thermococcus strains may be capable of virion production under appropriate conditions.

Several integrated elements show little similarity to any previously described MGE families, likely representing novel viral or plasmid lineages. These include a family of elements identified in *Py. sp.* IRI42c, *T. nautili* strains AMTc94, 29-3, 9-3, and *T. sp.* AMTc30 strain EXT11c (Fig 6), as well as singular MGEs such as those integrated in *T. sp.* IRI33c, *T. sp.* AMT11, and *Py. sp.* EXT15c.

The availability of multiple genomes per species further highlights the extent of strain-level mobilome variation. Among the 10 strains now classified as *T. nautili* (Table S3), the type strain 30-1^T^ carries the plasmids pTN1, pTN2, and pTN3 [106], whereas strain 33-3 carries the conjugative plasmid pT33-3 [62]. Beyond these well-characterized examples, additional strains harbor distinct complements of mobile elements, including a plasmid in *T. nautili* 9-3 and three co-existing plasmids in *T. nautili* E15P33. An assortment of pT26-2, pTN3, and TPV1 family integrated elements, as well as previously unclassified integrated MGEs are also distributed unevenly across *T. nautili* strains (Fig 6). Thus, mobilome composition varies extensively even within a genomically coherent species, such that any single reference genome captures only a fraction of the MGE diversity present in natural populations.

Collectively, these Thermococcales MGEs exhibit substantial diversity in size, gene content, and inferred lifestyle, with some element families confined to specific lineages, while others span multiple Thermococcales genera. These patterns establish a foundation for future work aimed at understanding how such elements infect, persist, and interact with their hosts.

## Discussion

Thermococcales are unique among archaeal lineages in that many hundreds of natural isolates are maintained in culture collections worldwide, and with our current dataset, 122 now have complete genome sequences. Unlike archaeal groups known primarily from metagenome-assembled genomes, Thermococcales combine dense genome sampling with experimentally tractable strains, enabling direct links between comparative genomics, physiology, and mobile element biology. Here, we combine order-wide pangenomic analysis and phylogenetic reconstructions to resolve taxonomic uncertainties and provide a genome-scale view of Thermococcales evolution.

Our work supports a revised genus-level classification within Thermococcales. In addition to the established genera *Pyrococcus, Palaeococcus,* and *Thermococcus,* we recover two additional monophyletic lineages previously classified as deeply branching *Thermococcus,* here proposed as “*Oleococcus*” and *“Piezococcus”.* Recognition of these clades resolves unstable phylogenetic placements observed under outdated taxonomies, and better reflects genome-scale evolutionary relationships. At the same time, the deeply branching *T. atlanticus* lineage likely represent additional genera; however, with only a single genome representative, we refrain from formally defining this without further sequences from related species.

Thermococcales genomes combine extreme structural plasticity with a strikingly constrained mode of functional diversification. While genome architectures are repeatedly reshuffled through inversions and rearrangements [24,28–31], variable functions, including metabolic pathways and mobile genetic elements, are gained and lost as intact multi-gene modules and preferentially localized to specific chromosomal regions (Fig. 6). This pattern contrasts with models of gradual, gene-by-gene metabolic evolution and instead points to a modular mode of genome evolution operating within strong organizational constraints, as previously suggested for essential gene clusters in Thermococcales [24].

Across the order, the core bioenergetic architecture is tightly conserved and forms part of the persistent pangenome. This conservation is coherent with decades of biochemical work on the archaeal “modified Emden-Meyerhof” pathway and its coupling to ferredoxin-based redox metabolism at high temperature. All Thermococcales encode archaea-specific glycolytic variants, including ADP-dependent kinases and ferredoxin-linked oxidation steps, together with sulfur reduction and hydrogenase systems that underpin anaerobic energy metabolism. These pathways show little evidence of gene loss or replacement across genera, indicating strong constraints on functions essential for growth in the hyperthermophilic anaerobic environments Thermococcales occupy.

In contrast, physiological surveys have shown that carbohydrate preferences and amino acid auxotrophies vary across Thermococcales species, often with weak phylogenetic signal. Our genome comparisons provide a mechanistic explanation – variation maps primarily to the gain and loss of intact multi-gene modules rather than incremental gene-level adaptation or erosion of pathways. Strikingly, homologous biosynthetic modules are shared across genera but are repeatedly inserted at genus-specific chromosomal locations. In *Pyrococcus,* variable amino acid biosynthesis pathways are embedded within a conserved locus flanking the 5S rRNA and tRNA^Asp^ genes, whereas closely related modules occupy a distinct region centered on the *folP-pheT* locus in *Piezococcus*. These observations indicate that module turnover is constrained not only by metabolic requirements but also by genome architecture. Rather than permitting unrestricted insertion across what appears to be highly scrambled chromosomes, Thermococcales genomes appear to contain a limited set of “permissive” integration loci that repeatedly accommodate large functional cassettes. Previous work demonstrated that the chromosomal positioning of essential gene clusters relative to the origin of replication is conserved across Thermococales despite pervasive inversions and rearrangements [24]. Our results extend this principle beyond the core genome, with even non-essential, horizontally acquired metabolic modules showing reproducible positional biases that are genus-specific. Genome architecture, therefore, constrains not only the conserved informational genes but also the integration of accessory functions. These constraints may reflect local features such as transcriptional context, replication timing, or higher-order chromosome structure, which influence whether newly acquired genes are functional and/or can be stably maintained alongside essential gene clusters. Identifying the determinants that define such permissive integration sites represents a testable direction for future work.

A similar architectural logic is seen in the distribution of integrated MGEs. All chromosomally integrated MGEs identified in this study encode an SSV1-like integrase that mediates site-specific recombination at select tRNA genes [29,32,33]. Consequently, viral and plasmid integration is inherently restricted to a defined set of conserved chromosomal loci. The same genomes that undergo extensive structural rearrangement therefore exhibit spatial constraint in both metabolic module acquisition and mobile element integration, as well as core gene modules.

At the same time, the composition of the mobilome remains highly dynamic. The extensive variation in MGE content observed among strains of *T. nautili* demonstrates that MGE repertoires can differ substantially even between closely related genomes. Thus while integration occurs at conserved chromosomal sites, the identity and combination of elements occupying those sites can shift rapidly. Single reference genomes therefore capture only a fraction of the diversity present in natural populations.

Taken together, these findings indicate that Thermococcales genomes accommodate rapid turnover of horizontally transferred modules within a limited set of conserved chromosomal loci, reconciling rapid functional diversification with long-term organizational consistency.

## Data availability

### Author contributions

RC and CL : Investigation, Data curation, Visualization, Writing – original draft. CB, ML, VS, DC: Investigation. EM,AW, MJ, PMO, LM, GE: Collection Resources. SF, GM, VB, JW : Sequencing Resources. PF : Funding acquisition, conceptualization, and project administration. JO: Supervision, Methodology. VDC : Conceptualization, Supervision, Methodology, Project administration, Writing – original draft. All authors: Read and approved the final manuscript.

### Funding and Acknowledgement

This work is supported by a European Research Council (ERC) grant from the European Union’s Seventh Framework Program (FP/2007-2013)/Project EVOMOBIL-ERC Grant agreement no. 340440. VDC and CL were supported by the French National Research Agency under the reference ANR-22-CPJ1-0082-01. We would like to thank the Microscope-Mage team and especially Zoé Rouy and Aurélie Genin-Lajus, and we would like to acknowledge “the LABGeM (CEA/Genoscope & CNRS UMR8030), the France Génomique and French Bioinformatics Institute national infrastructures (funded as part of Investissement d’Avenir program managed by Agence Nationale pour la Recherche, contracts ANR-10-INBS-09, ANR-11-INBS-0013 and ANR-21-ESRE-0048)” for their support within the MicroScope annotation platform.

## Supplementary Figures and Tables legends

**Sup Table 1:**
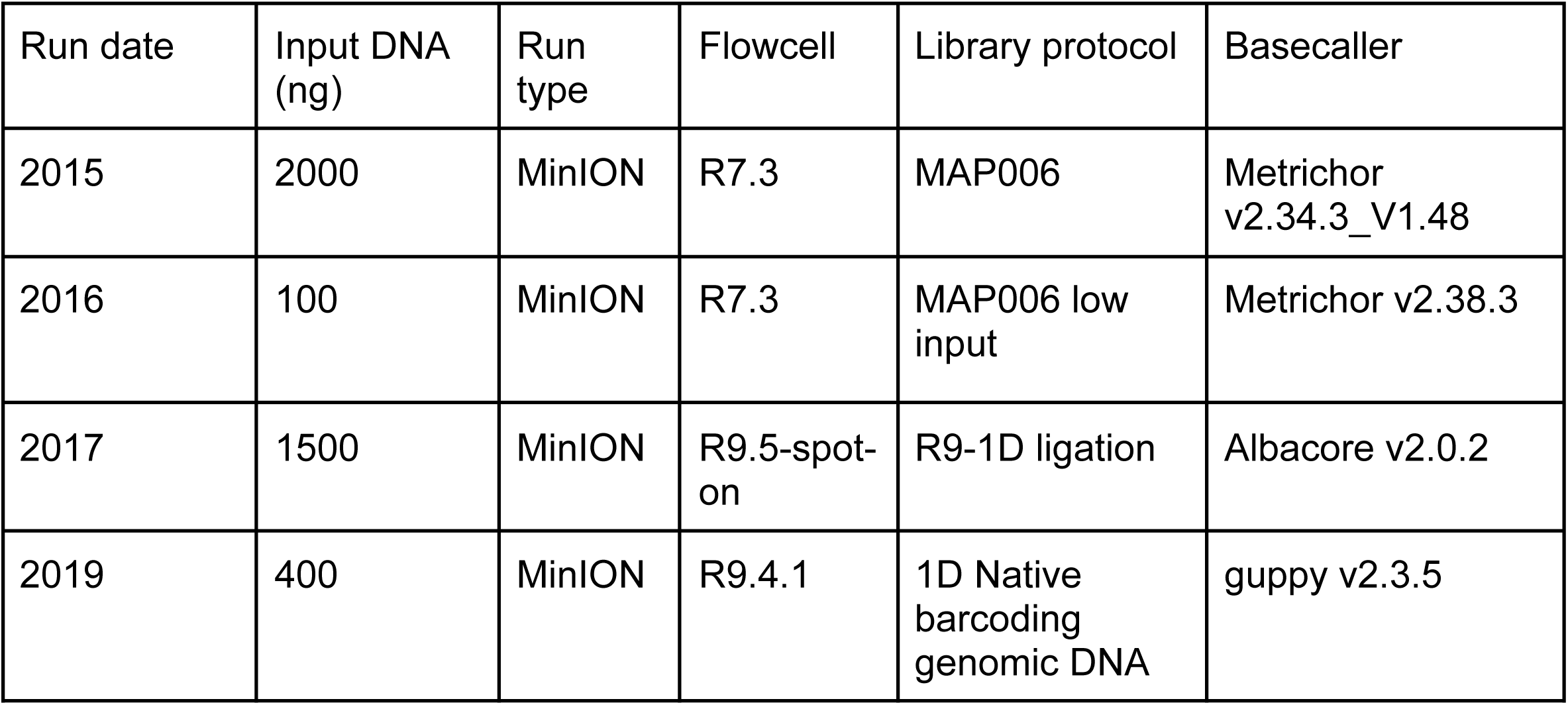
Oxford Nanopore Technologies used from 2015 to 2019 sequencing phases.

**Sup Table 2: Genomic characteristics, sampling metadata, and pangenome statistics of the analyzed Thermoccales genomes isolates**. For each strain of Thermococcales, the table provides the genome identifier, geographic origin (*Geographic Region*), sampling coordinates, sampling depth (*depth*), and genome features including genome size (*Genome Size, Mb*), GC content (*GC content, %*), chromosome completeness (*Complete chromosome*), protein-coding density (*% protein coding density*), average coding sequence length (*Average CDS length*), average intergenic region length (*Average intergenic length*), and the presence of extrachromosomal elements (*Extrachromosomal element*). Information on extrachromosomal genetic elements. Pangenome metrics comprise the total number of genes (*Genes*), gene families (*Families*), exact core families (*Exact core families*), persistent families (*Persistent families*), shell families (*Shell families*), and cloud families (*Cloud families*), together with their relative proportions (*% Persistent families*, *% Shell families*, *% Cloud families*, and *% Core families*).

**Sup Table 3: Taxonomic reassignment and species delineation of the analyzed Thermococcales genomes.** For each genome, the table reports the genome identifier (*Genome*), the previously assigned taxonomic name (*Previous Genus species name*), and the corresponding literature source (*Reference*). Updated taxonomic classifications are presented as the proposed genus (*Proposed Genus*) and species assignments inferred from Average Nucleotide Identity (*ANI-resolved species*), Mash distance (*Mash-resolved species*), and Average Amino Acid Identity (*AAI-resolved species*) analyses. The level of concordance among these taxonomic metrics is indicated by the number of agreeing methods (*Number of metrics agreeing*) and the agreement with the previously assigned species (*Agreement with previous species*). The final proposed taxonomic designation (*Proposed Genus species name*) and any relevant comments or considerations (*Notes*) are also provided.

**Sup Fig. 1 : Heatmap visualization of the diversity of the Thermococcales** - Heatmaps visualization of different evolutionary distance indicators with a phylogeny of the Thermococcale : A) Average nucleotide identity (ANI), only value greater than 77 are shown, the higher the value, the closer the genomes are. If the ANI between two genomes is above 76, they belong to the same genus, if it’s above 95, they belong to the same species (Jain et al. 2018). B) Amino acid identity (AAI) only the value greater than 65 are shown, the higher the value, the closer the genomes are. If the AAI between two genomes is above 75, they belong to the same genus, if it’s above 95, they belong to the same species (Jain et al. 2018) C) Mash distance only the value lesser than 0.3 are shown, the lower the value, the closer the genomes are. If the Mash distance between two genomes is below 0.3, they belong to the same genus, if it’s below 0.05, they belong to the same species (Ondov et al., 2016)

**Sup Fig. 2: Thermococcales Genomic and pangenomic data** - The phylogeny was performed with IQtree on the 122 Thermococcales genomes and an outgroup of 37 genomes from other group I Euryarchaeota with the 76 universal archaea markers of anvi’o. It has an alignment of 16433 positions and uses a LG+F+R10 model. The size of the genome is shown with a black histogram with a midpoint at 2 Mbp, the GC% is in red with a midpoint at 50%, and the histogram at the right of the figure is the repartition of the genes families for the 4 partitions of the pangenome (the core is in red, the persistent in orange, the shell in green and the cloud in blue).

**Sup Fig. 3 : Collapsed phylogenic tree of the Thermococcales built with different sets of76 archaeal core proteins** using Methanofastidiosales as an outgroup. - A) Collapsed phylogenic tree of non ribosomal proteins with an alignment of 10650 positions and the LG+F+R10 best model. B) Collapsed phylogenic tree of the ribosomal proteins with an alignment of 5319 positions and the LG+F+R4 best model. C) Collapsed phylogenic tree of the ribosomal proteins with an alignment of 16351 positions and the LG+F+R10 best model. For both phylogenies, nodes supported by an ultrafast bootstrap value >95 and an SH-aLRT support value >80 are marked with grey dots. Visualized with iTOL. In the different trees, the genomes belonging to the different genera defined in this analysis are indicated with different colors. The orange corresponds to Pyrococcus, the green to Oleococcus, the red to Palaeococcus, the blue to Piezococcus, and the purple to Thermococcus.

**Sup Fig. 4 : Phylogenetic tree of Thermococcales built with different the non ribosomal proteins using Methanofastidiosales as an outgroup.** - Collapsed phylogenic tree of non-ribosomal proteins with an alignment of 10650 positions and the LG+F+R10 best model. For all the trees, support is indicated by grey dots, which are the one with an ultrafast bootstrap support above 95 and an SH-like approximate likelihood ratio support above 80. Visualized with iTOL. In the tree, the genomes belonging to the different genera defined in this analysis are indicated with different colors. The orange corresponds to Pyrococcus, the green to Oleococcus, the red to Palaeococcus, the blue to Piezococcus and the purple to Thermococcus.

**Sup Fig. 5 : Phylogenetic tree of Thermococcales built with different the ribosomal proteins using Methanofastidiosales as an outgroup.** - Collapsed phylogenic tree of non-ribosomal proteins with an alignment of 5319 positions and the LG+F+R4 best model. For all the trees, support is indicated by grey dots, which are the one with an ultrafast bootstrap support above 95 and an SH-like approximate likelihood ratio support above 80. Visualized with iTOL. In the tree, the genomes belonging to the different genera defined in this analysis are indicated with different colors. The orange corresponds to Pyrococcus, the green to Oleococcus, the red to Palaeococcus, the blue to Piezococcus and the purple to Thermococcus.

**Sup Fig. 6 : Phylogenetic tree of Thermococcales built with the 76 archaeal core proteins using Methanofastidiosales as an outgroup.** - Collapsed phylogenic tree of non-ribosomal proteins with an alignment of 16351 positions and the LG+F+R10 best model. For all the trees, support is indicated by grey dots, if they meet the dual criteria of an ultrafast bootstrap support value higher than 95% and an SH-like approximate likelihood ratio support value greater than 80%. Visualized with iTOL. In the tree, the genomes belonging to the different genera defined in this analysis are indicated with different colors. The orange corresponds to Pyrococcus, the green to Oleococcus, the red to Palaeococcus, the blue to Piezococcus and the purple to Thermococcus.

**Sup Fig. 7: Examples of individual gene trees and rogue taxa identification.** Here are four single-gene phylogenetic trees reconstructed from four core proteins (HypE, HypF, Slipinfamily protein, and UbiD family decarboxylase). Rogue taxa were detected based on their inconsistent placements across tree reconstructions and are highlighted in the figure. These taxa were subsequently considered for exclusion to improve phylogenetic resolution and node support in downstream analyses. In the trees, the genomes belonging to the different genera defined in this analysis are indicated with different colors. The orange corresponds to Pyrococcus, the green to Oleococcus, the red to Palaeococcus, the blue to Piezococcus and the purple to Thermococcus.

**Sup Fig. 8: Phylogenic tree of the 122 Thermococcales genomes from the concatenation of 142 core protein without the identified rogues taxa**. The tree was performed with an alignment of 39788 positions by using a partition approach with the best model determined before for each individual proteins. Nodes represented by blue points meet the dual criteria of an ultrafast bootstrap support value higher than 95% and an SH-like approximate likelihood ratio support value greater than 80%. Visualized with iTOL. In the tree, the genomes belonging to the different genera defined in this analysis are indicated with different colors. The orange corresponds to Pyrococcus, the green to Oleococcus, the red to Palaeococcus, the blue to Piezococcus and the purple to Thermococcus.

**Sup Fig. 9: Phylogenetic tree of the concatenation of the nucleoside ABC transporter proteins, a horizontal gene transfer example.** The nucleoside ABC transporter (K23537, K23535, K23536) was predicted using KOfam-based annotation. The corresponding *Pyrococcus* proteins (PF1696-PF1698) exhibited higher sequence identity to homologous colocalized genes found in genomes or metagenome-assembled genomes (MAGs) of hydrothermal vent-associated Crenarchaeota (52-61%), such as *Staphylothermus marinus* and *Candidatus Tiamatella incendiivivens* [109,110] (indicated in pink in the tree), than to the other Thermococales genomes (35-42%) with the classical colors schema. The orange correspond to Pyrococcus, the green to Oleococcus, the red to Palaeococcus, the blue to Piezococcus, and the purple to Thermococcus. Nodes represented by blue points meet the dual criteria of an ultrafast bootstrap support value higher than 95% and an SH-like approximate likelihood ratio support value greater than 80%.

## Notes

### Competing Interest Statement

The authors have declared no competing interest.

