## Supplementary table 1 for "Thermococcales pangenome uncovers new genera and modular evolution"

**Table S1**: Oxford Nanopore Technologies used from 2015 to 2019 sequencing phases.

| Run date | Input DNA (ng) | Run type | Flowcell | Library protocol | Basecaller |
|---|---|---|---|---|---|
| 2015 | 2000 | MinION | R7.3 | MAP006 | Metrichor v2.34.3_V1.48 |
| 2016 | 100 | MinION | R7.3 | MAP006 low input | Metrichor v2.38.3 |
| 2017 | 1500 | MinION | R9.5-spot-on | R9-1D ligation | Albacore v2.0.2 |
| 2019 | 400 | MinION | R9.4.1 | 1D Native barcoding genomic DNA | guppy v2.3.5 |
