## Supplementary table 2 for "Thermococcales pangenome uncovers new genera and modular evolution"

| Name | Genome | Assembly accessic | Geographic Regior | Geographic_locati | *Geographic_locat* | sample | depth | Collection_date | Genome_Size_Mb | GC_content% | Complete_chromo | % protein codingD | Average_CDS_leng | Average_intergen | Extrachromosoma | EGM_name | acces_EGM | Genes | Families | Exact_core_famili | Persistent_familie | Shell_families | Cloud_families | %_Persistent_fam | %_Shell_families | %_Cloud_families | %_Core_families |
|---|---|---|---|---|---|---|---|---|---|---|---|---|---|---|---|---|---|---|---|---|---|---|---|---|---|---|---|
| Palaeococcus_pacificus_DY20341 | CP006019 | GCA_000725425,1 | East Pacific Ocean | -1,616667 | -102,75 | hydrothermal vent | 2,737 m | 2008 | 1.859370 | 43.04 | yes | 91.43 | 835.67 | 92.44 | | | | 1969 | 1952 | 483 | 1111 | 371 | 470 | 56.91598361 | 19.00614754 | 24.07786885 | 24.74385246 |
| Thermococcus_indicus_IOH1 | NZ_CP040846 | GCA_006274605,1 | Indian Ocean: Onn | -11,24 | 66,25 | hydrothermal vent | 2,021m | 2018 | 2.2 | 55 | yes | 88.3 | 842.67 | 107.52 | 1 | pTI1 | NZ_CP040847,1 | 2357 | 2305 | 483 | 1257 | 725 | 323 | 54.53362256 | 31.45336226 | 14.01301518 | 20.95444685 |
| Thermococcus_argininiproducens_IOH2 | NZ_CP080572,1 | GCA_023746595,1 | Central Indian Oce | -11,4 | 66,91 | sediment | 2018 | 2018 | 1.94625 | 39.4 | yes | 90.86 | 839 | 96.71 | | | | 2063 | 2020 | 483 | 1102 | 439 | 479 | 54.55445545 | 21.73267327 | 23.71287129 | 23.91089109 |
| Pyrococcus_abyssi_GE5 | NC_000868 | GCA_000195935,2 | North Fiji Basin of | -16,5 | 173,5 | Active chimney in | 3,500m | 1989 | 1.765118 | 44.5 | yes | 92.91 | 863.61 | 99.5 | 1 | pGT5 | NC_001773,1/ | 1892 | 1867 | 483 | 1031 | 499 | 337 | 55.22228174 | 26.72737011 | 18.05034815 | 25.87038029 |
| Pyrococcus_abyssi_GE2 | 7482 | PABYGE2,1 | North Fiji Basin of | -16,5 | 173,5 | Active chimney in | 3,500m | | 1.815336 | 44.56 | yes | 92.46 | 848.17 | 91.26 | | | | 1987 | 1954 | 483 | 1019 | 490 | 445 | 52.14943705 | 25.07676561 | 22.77379734 | 24.7185261 |
| Pyrococcus_sp_GE_23 | 8812 | PGE23,1 | North Fiji Basin of | -16,5 | 173,5 | Active chimney in | 3,500m | | 1.720484 | 44.72 | yes | 92.68 | 841.53 | 84.84 | 1 | plasmid PGE23_p,1 | PGE23_p,1 | 1904 | 1872 | 483 | 1020 | 466 | 386 | 54.48717949 | 24.89316239 | 20.61965812 | 25.80128205 |
| Thermococcus_fumicolans_ST557 | 14021 | Tfumi_Ch,1 | the White Lady hy | -16,9833 | -173 | deep-sea hydrothe | 2000 | 1989 | 1.961384 | 55.53 | yes | 92.04 | 802.01 | 85.34 | 1 | Tfumi_p,1 | | 2260 | 2188 | 483 | 1221 | 446 | 521 | 55.80438757 | 20.38391225 | 23.81170018 | 22.0749543 |
| Thermococcus_sp_690 | | | Pacific Ocean, Lau | -20,066251 | -177,041931 | Marine hydrotherm | 2,100m | 1989 | 1.933537 | 55.54 | yes | 92.07 | 825.34 | 86.56 | 2 | TH690pa,1; TH690pb,1 | | 2121 | 2091 | 483 | 1234 | 426 | 431 | 59.01482544 | 20.37302726 | 20.6121473 | 23.0989957 |
| Thermococcus_barophilus_423_3_23 | 4061 | Tb423_Chr,1 | Pacific Ocean, Lau | -20,066251 | -177,041931 | Marine hydrotherm | 2,100m | | 1.974483 | 41.67 | yes | 92.09 | 822.89 | 88.48 | | | | 2217 | 2170 | 483 | 1152 | 485 | 533 | 53.0875576 | 22.35023041 | 24.56221198 | 22.25806452 |
| Thermococcus_barophilus_DT4 | 4060 | TbDT4_Chr,1 | Pacific Ocean, Lau | -20,066251 | -177,041931 | Marine hydrotherm | 2,100m | | 2.098682 | 40.37 | yes | 91.89 | 808.66 | 90.97 | | | | 2401 | 2310 | 483 | 1129 | 447 | 734 | 48.87445887 | 19.35064935 | 31.77489177 | 20.90909091 |
| Thermococcus_onnurineus_NA1 | CP000855 | GCA_000018365,1 | PACManus (Papua-/ | -3,721111 | 151,684167 | hydrothermal vent | 1,650m | 2002 | 1.84761 | 51.3 | yes | 90.65 | 831.04 | 103 | | | | 1979 | 1938 | 483 | 1227 | 465 | 246 | 63.3126935 | 23.99380805 | 12.69349845 | 24.92260062 |
| Pyrococcus_sp_NA2 | CP002670 | GCA_000211475,1 | PACManus | -3,721111 | 151,684167 | hydrothermal vent | 1,650m | 2002 | 1.86132 | 42.7 | yes | 92.02 | 837.7 | 101.2 | 0 | | | 1968 | 1938 | 483 | 1009 | 499 | 430 | 52.06398349 | 25.74819401 | 22.1878225 | 24.92260062 |
| Thermococcus_gorgonarius_W_12 | CP014855 | GCA_002214385,1 | Bay of Plenty, New | -36,88 | 176,38 | sand from the shore of Whale Isalnd, i | | 1990 | 1.674122 | 51.5 | yes | 89.82 | 824.4 | 91.81 | | | | 1841 | 1820 | 483 | 1160 | 306 | 354 | 63.73626374 | 16.81318681 | 19.45054945 | 26.53846154 |
| Thermococcus_pacificus_P4 | CP015102 | GCA_002214485,1 | Bay of Plenty, New | -37,62 | 177,14 | bottom deposits of | 40m | 1990 | 1.78567 | 54.24 | yes | 90.81 | 825.6 | 101.97 | | | | 1925 | 1901 | 483 | 1240 | 410 | 251 | 65.22882693 | 21.567596 | 13.20357706 | 25.40768017 |
| Thermococcus_aciditolerans_SY113 | NZ_CP041932,1 | GCA_008152015,1 | Indian Ocean | -37,78 | -49,65 | hydrothermal vent | 2770 m | 2018 | 2.2 | 54.5 | yes | 90.21 | 835.54 | 100.46 | | | | 2333 | 2280 | 483 | 1259 | 685 | 336 | 55.21929825 | 30.04385965 | 14.73684211 | 21.18421053 |
| Thermococcus_sp_SY098 | | GCA_035621495 | marine hydrotherm | -37,78 | 49,65 | | 2723 | 2018 | 2.033621 | 41.5 | yes | 91.96 | 818.97 | 88.21 | | | | 2234 | 2211 | 483 | 1152 | 507 | 552 | 52.10312076 | 22.93080054 | 24.9660787 | 21.84531886 |
| Thermococcus_thermotolerans_813A4 | NZ_CP102602,1 | GCA_024707485,1 | LongQi hydrotherm | -37,7827949 | 49,6497126 | chimney in the Sou | 2,739m | 2019 | 2.07086 | 53.5 | yes | 91.82 | 826.19 | 89.96 | | | | 2280 | 2240 | 483 | 1261 | 747 | 232 | 56.29464286 | 33.34821429 | 10.35714286 | 21.5625 |
| Thermococcus_zilligii_AN1 | | GCA_000258515,1 | Rotorua, Kuirau Pa | -38,13 | 176,25 | anaerobic mud from alkaline hot sprir | NA | | 1.76456 | 54.5 | no | 90.42 | 843.02 | 105.1 | | | | 1817 | 1796 | 483 | 1147 | 327 | 322 | 63.86414254 | 18.20712695 | 17.92873051 | 26.89309577 |
| Thermococcus_waiotapuensis_WT1 | 14024 | | New Zealand | -38,7608 | 175,91453 | Waiotapu hot spring, Lake Taupo area | NA | | 1.801659 | 53.46 | yes | 97.09 | 817.81 | 97.09 | | | | 2012 | 1942 | 483 | 1149 | 358 | 435 | 59.16580844 | 18.4346035 | 22.39958805 | 24.87126674 |
| Thermococcus_prieuri_Col3 | 7494 | TPCOL3,1 | East Pacific Rise | -7,4233333 | -107,8016667 | hydrothermal chim | 2700 m | 2004 | 1.927526 | 54.27 | yes | 92.15 | 830.33 | 91.88 | | | | 2150 | 2134 | 483 | 1250 | 453 | 431 | 58.57544517 | 21.22774133 | 20.1968135 | 22.63355201 |
| Thermococcus_sp_TPV | 7510 | | East Pacific Rise | -7,4233333 | -107,8016667 | hydrothermal chim | 2700 m | 2004 | 2.114736 | 54.53 | yes | 91.77 | 839.89 | 98.21 | | | | 2319 | 2281 | 483 | 1238 | 712 | 331 | 54.27444103 | 31.21437966 | 14.51117931 | 21.17492328 |
| Thermococcus_sp_AM4 | CP002952 | GCA_000151205,2 | East Pacific Rise | 12,13333 | -103,93333 | hydrothermal vent | 2,600m | a2011 | 2.086428 | 54.8 | yes | 91.17 | 832 | 96.28 | | | | 2256 | 2229 | 483 | 1195 | 502 | 532 | 53.61148497 | 22.52131 | 23.86720502 | 21.66890983 |
| Thermococcus_33N | MAGE A recuperer | MAGE A recuperer | East Pacific Ridge, | 12,7944444 | -104,0388889 | hydrothermal chim | 2,630m | 1999 | 1.916721 | 54.21 | yes | 91.78 | 866.58 | 92.98 | | | | 2039 | 2022 | 483 | 1277 | 592 | 153 | 63.15529179 | 29.27794263 | 7.566765579 | 23.88724036 |
| Pyrococcus_31_3 | 7490 | PYRO313_b,1 | East Pacific Ridge, | 12,811382 | -103,94059 | hydrothermal chim | 2,632m | 1999 | 1.989130 | 44.97 | yes | 91.95 | 822.14 | 98.87 | | | | 2242 | 2191 | 483 | 1046 | 625 | 520 | 47.74075764 | 28.52578731 | 23.73345504 | 22.04472843 |
| Pyrococcus_32_1 | 7495 | PYRO321,1 | East Pacific Ridge, | 12,811382 | -103,94059 | hydrothermal chim | 2,632m | 1999 | 1.989216 | 44.97 | yes | 92 | 822.53 | 98.54 | 3 | | | 2241 | 2190 | 483 | 1045 | 626 | 519 | 47.71689498 | 28.58447489 | 23.69863014 | 22.05479452 |
| Pyrococcus_sp_32_3 | 8815 | PYRO323,1 | East Pacific Ridge, | 12,811382 | -103,94059 | hydrothermal chim | 2,632m | 1999 | 1.989130 | 44.97 | yes | 91.94 | 820.96 | 98.88 | 3 | | | 2245 | 2195 | 483 | 1046 | 625 | 524 | 47.65375854 | 28.4738041 | 23.87243736 | 22.00455581 |
| Pyrococcus_sp_32_4 | 8823 | PYRO324,1 | East Pacific Ridge, | 12,811382 | -103,94059 | hydrothermal chim | 2,632m | 1999 | 1.989130 | 44.97 | yes | 92.04 | 820.94 | 97.83 | 1 | | | 2246 | 2195 | 483 | 1045 | 626 | 524 | 47.60820046 | 28.51936219 | 23.87243736 | 22.00455581 |
| Thermococcus_AMTc30 | 10769 | TAMTC30,1 | East Pacific Ridge, | 12,811382 | -103,94059 | hydrothermal chim | 2,632m | 1999 | 1.941830 | 54.89 | yes | 92.18 | 839.29 | 93.05 | | | | 2141 | 2114 | 483 | 1251 | 662 | 201 | 59.1769158 | 31.31504257 | 9.508041627 | 22.84768212 |
| Thermococcus_sp_AMTc71 | 13480 | TAMTC71,2 | East Pacific Ridge, | 12,811382 | -103,94059 | hydrothermal chimney | | 1999 | 1.954161 | 55.96 | yes | 92 | 849.22 | 89.98 | | | | 2123 | 2103 | 483 | 1249 | 582 | 272 | 59.3913457 | 27.67475036 | 12.93390395 | 22.96718973 |
| Thermococcus_sp_AMTc72 | 13481 | TAMTC72,2 | East Pacific Ridge, | 12,811382 | -103,94059 | hydrothermal chimney | | 1999 | 1.954162 | 55.96 | yes | 92.04 | 848.41 | 89.54 | | | | 2126 | 2106 | 483 | 1249 | 583 | 274 | 59.30674264 | 27.68281102 | 13.01044634 | 22.93447293 |
| Thermococcus_31_3 | 7489 | THER313_a,1 | East Pacific Ridge, | 12,811382 | -103,94059 | hydrothermal chim | 2,632m | 1999 | 2.026285 | 54.86 | yes | 92.32 | 862.15 | 89.01 | | | | 2178 | 2157 | 483 | 1270 | 580 | 307 | 58.8780714 | 26.88919796 | 14.23273064 | 22.3922114 |
| Thermococcus_sp_AMTc102 | 11786 | THAMT102,1 | East Pacific Rise, T | 12,812438 | -103,940327 | hydrothermal chimney extremity | | 1999 | 2.040569 | 55.79 | yes | 92.45 | 823.59 | 90.46 | | | | 2307 | 2260 | 483 | 1220 | 688 | 352 | 53.98230088 | 30.44247788 | 15.57522124 | 21.37168142 |
| Thermococcus_sp_26_2 | GCA_ | THER262,1 | East Pacific Rise, T | 12,812438 | -103,940327 | | | 1999 | 2.030483 | 55.89 | yes | 92.47 | 820.58 | 91.23 | | | | 2300 | 2252 | 483 | 1224 | 697 | 331 | 54.35168739 | 30.95026643 | 14.69804618 | 21.44760213 |
| Thermococcus_23_2 | 7487 | THER232,1 | East Pacific Ridge, | 12,813095 | -103,939486 | hydrothermal chim | 2,610m | 1999 | 1.854909 | 55.69 | yes | 92.44 | 856.51 | 84.65 | | | | 2012 | 1995 | 483 | 1279 | 535 | 181 | 64.11027569 | 26.81704261 | 9.072681704 | 24.21052632 |
| Thermococcus_sp_23_4 | 9793 | THER234,1 | East Pacific Rise, C | 12,813095 | -103,939486 | hydrothermal chim | 2,610m | 1999 | 1.854775 | 55.69 | yes | 92.43 | 857.4 | 83.38 | | | | 2008 | 1990 | 483 | 1279 | 533 | 178 | 64.27135678 | 26.7839196 | 8.944723618 | 24.27135678 |
| Thermococcus_sp_9_3 | 9794 | THER93,1 | East Pacific Rise, C | 12,813095 | -103,939486 | hydrothermal chim | 2,610m | 1999 | 2.021135 | 54.85 | yes | 92.13 | 825.18 | 91.49 | 1 | plasmid THER93_p,1 | | 2264 | 2230 | 483 | 1258 | 790 | 182 | 56.41255605 | 35.42600897 | 8.161434978 | 21.65919283 |
| Thermococcus_sp_AMTc09 | 12160 | TAMTC09,1 | East Pacific Ridge, | 12,813149 | -103,93932 | | | 1999 | 2.041610 | 55.82 | yes | 92.36 | 815.25 | 90.37 | | | | 2332 | 2289 | 483 | 1216 | 699 | 374 | 53.12363478 | 30.53735256 | 16.33901267 | 21.10091743 |
| Thermococcus_nautili_30_1 | CP007264 | GCA_000585495,1 | East Pacific Rise | 12,831631 | -103,94654 | | | 1999 | 2 | 55 | yes | 91.74 | 829.89 | 96.28 | 3 | pTN1,pTN2,pTN3 | | 2168 | 2139 | 483 | 1259 | 746 | 134 | 58.85928004 | 34.87611033 | 6.264609631 | 22.58064516 |
| Thermococcus_sp_5_4 | CP021848 | GCA_002197185,1 | East Pacific Ocean | 12,831631 | -103,94654 | deep-sea hydrothe | 2,330m | 1999 | 1.85491 | 55.7 | yes | 92.45 | 853.76 | 82.62 | | | | 1975 | 1958 | 483 | 1284 | 536 | 138 | 65.57711951 | 27.37487232 | 7.048008172 | 24.6680286 |
| Thermococcus_AMTc11 | 7483 | TAMTC11,1 | East Pacific Ridge, | 12,831658 | -103,946531 | hydrothermal chim | 2,634m | 1999 | 2.378328 | 54.47 | yes | 91.32 | 828.85 | 100.48 | | | | 2628 | 2564 | 483 | 1278 | 784 | 502 | 49.84399376 | 30.57722309 | 19.57878315 | 18.83775351 |
| Thermococcus_sp_AMTc79 | 11778 | TAMTC79,1 | East Pacific Ridge, | 12,831658 | -103,946531 | | | 1999 | 1.977274 | 54.52 | yes | 91.86 | 817.82 | 93.65 | | | | 2234 | 2203 | 483 | 1257 | 718 | 228 | 57.05855651 | 32.59192011 | 10.34952338 | 21.92464821 |
| Thermococcus_AMTc85 | 10774 | TAMTC85,1 | East Pacific Ridge, | 12,831658 | -103,946531 | hydrothermal source, colonization mo | | 1999 | 1.933709 | 54.48 | yes | 92.06 | 824.15 | 90.12 | | | | 2169 | 2146 | 483 | 1248 | 634 | 264 | 58.15470643 | 29.54333644 | 12.30195713 | 22.50698975 |
| Thermococcus_29_3 | 7513 | THER293,1 | East Pacific Ridge, | 12,831658 | -103,946531 | hydrothermal chim | 2,634m | 1999 | 1.969639 | 54.8 | yes | 91.9 | 836.42 | 96.7 | | | | 2173 | 2142 | 483 | 1242 | 758 | 142 | 57.98319328 | 35.38748833 | 6.629318394 | 22.54901961 |
| Thermococcus_nautili_33_3 | 8825 | THER333,1 | East Pacific Rise, F | 12,831658 | -103,946531 | hydrothermal chim | 2,634m | 1999 | 2.013327 | 54.6 | yes | 91.94 | 816.91 | 93.19 | | | | 2272 | 2235 | 483 | 1258 | 746 | 231 | 56.28635347 | 33.37807606 | 10.33557047 | 21.61073826 |
| Thermococcus_AMTc51 | 10770 | TAMTC51,1 | East Pacific Ridge, | 12,83947 | -103,948737 | hydrothermal chim | 2,635m | 1999 | 1.941015 | 53.93 | yes | 91.57 | 843.14 | 100.92 | | | | 2115 | 2088 | 483 | 1239 | 579 | 270 | 59.33908046 | 27.72988506 | 12.93103448 | 23.13218391 |
| Thermococcus_sp_AMTc94 | 11802 | TAMTC94,1 | East Pacific Rise, L | 12,83947 | -103,948737 | hydrothermal chim | 2,635m | 1999 | 1.968088 | 55 | yes | 92.57 | 827.17 | 89.56 | | | | 2212 | 2182 | 483 | 1254 | 769 | 159 | 57.47021082 | 35.24289643 | 7.286892759 | 22.13565536 |
| Thermococcus_komboloii_15_2 | 7488 | THER152,1 | East Pacific Ridge, | 12,83947 | -103,948737 | hydrothermal chim | 2,635m | 1999 | 1.881790 | 54.44 | yes | 92.08 | 845.88 | 91.73 | | | | 2060 | 2044 | 483 | 1248 | 446 | 350 | 61.05675147 | 21.81996086 | 17.12328767 | 23.63013699 |
| Thermococcus_barophilus_CH1 | 4059 | TbCH1_Chr,1 | Atlantic North, Ash | 12,970000 | -44,86 | hydrpthermal vent | 4,100m | | 2.171602 | 41.73 | yes | 92.27 | 836.96 | 91.35 | | | | 2408 | 2356 | 483 | 1142 | 525 | 689 | 48.47198642 | 22.28353141 | 29.24448217 | 20.5008489 |
| Pyrococcus_yayanosii_CH1 | CP002779 | GCA_000215995,1 | Atlantic Ocean, mi | 12,9733 | -44,8633 | deep sea hydrothe | 4,100m | 2007 | 1.71682 | 51.6 | yes | 90.59 | 812.94 | 89.49 | 0 | | | 1857 | 1821 | 483 | 1036 | 390 | 395 | 56.89181768 | 21.41680395 | 21.69137836 | 26.52388797 |
| Thermococcus_barophilus_CH5 | CP013050 | GCA_001433455,1 | Atlantic Ocean, mi | 14,75 | -44,98 | hydrothermal fluid | 3,500m | 1993 | 2.38853 | 41.8 | yes | 91.59 | 809.44 | 95.97 | 0 | | | 2589 | 2505 | 483 | 1140 | 605 | 760 | 45.50898204 | 24.15169661 | 30.33932136 | 19.28143713 |
| Thermococcus_piezophilus_CDGS | NZ_CP015520 | GCA_001647085,1 | Mexico Beebe vent | 18,32 | -81,43 | hydrothermal vent | | 2010 | 1.9 | 51 | yes | 80.57 | 757.05 | 116.27 | | | | 2108 | 2021 | 483 | 1176 | 445 | 400 | 58.18901534 | 22.01880257 | 19.79218209 | 23.89905987 |
| Thermococcus_chitonophagus_isolate_1 | LN999010 | GCA_900012635,1 | hydrothermal vent | 20,50 | -109,06 | hydrothermal vent | | 1991 | 1.969648 | 45 | yes | 90.24 | 806.45 | 96.42 | | | | 2141 | 2104 | 483 | 1040 | 586 | 478 | 49.42965779 | 27.85171103 | 22.71863118 | 22.95627376 |
| Thermococcus_chitonophagus | CP015193 | GCA_002214605,1 | Mexico: Off the me | 20,83 | -109,1 | hydrothermal fluid | 2600 | 1991 | 2 | 45 | yes | 90.24 | 806.45 | 96.42 | | | | 2127 | 2095 | 483 | 1036 | 585 | 474 | 49.45107399 | 27.92362768 | 22.62529833 | 23.0548926 |
| Thermococcus_sp_BubbBath | SAMN10963467 | GCA_012027595,1 | Mariana Arc | 21,19505 | 144,11488 | megaplume | | 1988-2008 | 2.0488 | 52.2 | no | 89.88 | 793.58 | 112.45 | | | | 2115 | 2087 | 483 | 1111 | 392 | 584 | 53.23430762 | 18.78294202 | 27.98275036 | 23.14326785 |
| Thermococcus_marinus_DSM_15227 | 11784 | 11784 | Atlantic Ocean Mid | 23,22 | -44,57 | | | 1995 | 2.078542 | 53.74 | yes | 90.83 | 824.3 | 104.68 | | | | 2300 | 2248 | 483 | 1235 | 635 | 378 | 54.93772242 | 28.24733096 | 16.81494662 | 21.48576512 |
| Thermococcus_barophilus_MP | NC_014804,1 | GCA_000151105,2 | Mid-Atlantic Ridge | 23,224118 | -44,564984 | hydrothermal vent | 3,550m | 1993 | 2.06424 | 41.71 | yes | 92.42 | 815.62 | 87.46 | 1 | pTBMP1 | | 2213 | 2182 | 483 | 1143 | 507 | 532 | 52.38313474 | 23.2355637 | 24.38130156 | 22.13565536 |
| Thermococcus_atlanticus_DSM_15226 | 11783 | THATL,1 | Mid Atlantic Ridge, | 26,133333 | -44,816667 | hydrothermal samp | 3,670m | a2002 | 2.294590 | 50.63 | yes | 91.26 | 844.28 | 104.52 | 1 | plasmid THATL_p,1 | | 2493 | 2434 | 483 | 1181 | 613 | 640 | 48.52095316 | 25.18488085 | 26.29416598 | 19.84387839 |
| Thermococcus_eurythermalis_A501 | NZ_CP008887 | GCA_000769655,1 | Pacific Ocean: Gul | 27,0067 | -110,409 | oil-immersed samp | 2006,9 | 2009 | 2.1 | 53.5 | yes | 89.82 | 841.86 | 108.32 | | | | 2250 | 2210 | 483 | 1233 | 634 | 343 | 55.7918552 | 28.68778281 | 15.52036199 | 21.85520362 |
| Pyrococcus_kukulkanii_NCB100 | CP010835 | GCA_001577775,1 | Guaymas Basin , G | 27,010567 | -111,40675 | Rebecca's Roost ve | 1,997m | 2010 | 1.97713 | 44.6 | yes | 86.23 | 805.41 | 104.54 | | | | 2153 | 2106 | 483 | 1033 | 567 | 506 | 49.05033238 | 26.92307692 | 24.02659069 | 22.93447293 |
| Thermococcus_gammatolerans_EJ3 | CP001398 | GCA_000022365,1 | Guaymas Basin | 27,0167 | -111,4 | hydrothermal chim | 2,616m | 1991 | 2 | 53.5 | yes | 92.98 | 873.41 | 83.99 | | | | 2167 | 2138 | 483 | 1166 | 480 | 492 | 54.53695042 | 22.45088868 | 23.0121609 | 22.59120674 |
| Thermococcus_radiotolerans_EJ2 | CP015106 | GCA_002214565,1 | Guayma Basin | 27,0167 | -111,4 | chimney of deep-s | 2,616m | 1991 | 1.868990 | 55.6 | yes | 92.32 | 845.72 | 85.81 | | | | 2004 | 1988 | 483 | 1269 | 570 | 149 | 63.83299799 | 28.67203219 | 7.494969819 | 24.29577465 |
| Thermococcus_sp_4557 | NC_015865 | GCA_000221185,1 | Gulf of California, | 27,25 | -111,5 | deep-sea hydrothe | 2,000m | a2011 | 2.01132 | 56.1 | yes | 91.86 | 845.65 | 93.44 | | | | 2112 | 2091 | 483 | 1253 | 635 | 203 | 59.92348159 | 30.36824486 | 9.708273553 | 23.0989957 |
| Thermococcus_guaymasensis_DSM_11113 | CP007140 | GCA_000816105,1 | Gulf of California, | 27,25 | -111,5 | hydrothermal vent | 2,000m | a1998 | 1.9 | 53 | yes | 90.87 | 816.99 | 95.09 | | | | 2077 | 2051 | 483 | 1186 | 433 | 432 | 57.825451 | 21.11165285 | 21.06289615 | 23.54948805 |
| Thermococcus_aggregans_DSM_12819 | 11781 | THAGGR,1 | Gulf of California, | 27,25 | -111,5 | | 2,000 m | a1999 | 1.890956 | 43.35 | yes | 90.98 | 828.1 | 98.66 | | | | 2085 | 2034 | 483 | 1093 | 433 | 508 | 53.73647984 | 21.28810226 | 24.9754179 | 23.74631268 |
| Thermococcus_aggregans_TY | NZ_CP099582,1 | GCA_024022995,1 | Mexico: Guaymas E | 27,51 | -100,57 | hydrothermal vent | | 1999 | 1.9 | 43 | yes | 90.62 | 827.76 | 97.65 | | | | 2000 | 1960 | 483 | 1092 | 431 | 437 | 55.71428571 | 21.98979592 | 22.29591837 | 24.64285714 |
| Thermococcus_siculi_RG_20 | CP015103 | GCA_002214505,1 | Mid-Okinawa Troug | 27,545 | 126,975 | deep-sea thermal | 1,394m | 1996 | 2.02917 | 55 | yes | 90.9 | 855.04 | 97.83 | | | | 2129 | 2110 | 483 | 1255 | 590 | 265 | 59.47867299 | 27.96208531 | 12.55924171 | 22.89099526 |
| Thermococcus_profundus_DT_5432 | NZ_CP014862 | GCA_002214585,1 | Middle Okinawa Tr | 27,55 | 126,966 | deep sea hydrothe | 1,395m | a1995 | 2.005098 | 53.125 | yes | 90.5 | 868.91 | 101.28 | 1 | unnamed | CP014863,1 | 2111 | 2084 | 483 | 1207 | 425 | 452 | 57.91746641 | 20.39347409 | 21.6890595 | 23.17658349 |
| Thermococcus_profundus_DSM_9503 | 4062 | TPro_Chr,1 | Pacific Ocean, Mid | 27,55 | 126,966 | deep sea hydrothe | 1,395m | a1994 | 2.005099 | 53.26 | yes | 91.7 | 877.28 | 98.21 | 1 | plasmid TPro_Plasmid,1 | | 2101 | 2072 | 483 | 1204 | 417 | 451 | 58.10810811 | 20.12548263 | 21.76640927 | 23.31081081 |
| Pyrococcus_horikoshii_OT3 | BA000001 | GCA_000011105,1 | Pacific Ocean, Oki | 27,5500 | 126,933 | marine hydrotherm | 1,395m | a1999 | 1.7385 | 41.9 | yes | 91.61 | 823.48 | 109.75 | | | | 1850 | 1829 | 483 | 987 | 446 | 396 | 53.96391471 | 24.38490979 | 21.65117551 | 26.40787315 |
| Thermococcus_celericrescens_DSM_17994 | SAMN04166689 | GCA_001484195,1 | Japan Western Pac | 28,34 | 140,38 | hydrothermal vent | | 2001 | 2.337139 | 54.5 | No | 90.04 | 794.41 | 115.55 | | | | 2412 | 2335 | 483 | 1212 | 720 | 403 | 51.90578158 | 30.83511777 | 17.25910064 | 20.68522484 |
| Thermococcus_peptonophilus_OG_1 | NZ_CP014750 | GCA_001592435,1 | Pacific Ocean Izu-E | 28,57 | 140,63 | Marine hydrotherm | 1380 | a1995 | 1.9 | 51.5 | yes | 89.53 | 837.03 | 102.67 | 1 | | CP014751,1 | 2024 | 2001 | 483 | 1239 | 428 | 334 | 61.91904048 | 21.38930535 | 16.69165417 | 24.13793103 |
| Thermococcus_coalescens_TS1 | 14020 | Tcoa_Ch1,1 | Pacific Ocean, Izu- | 28,5714 | 140,644 | Hydrothermal fluic | 1380 | 2001 | 2.134407 | 54.26 | yes | 91.46 | 834.21 | 97.35 | | | | 2347 | 2306 | 483 | 1257 | 706 | 343 | 54.50997398 | 30.61578491 | 14.87424111 | 20.94535993 |
| Thermococcus_kodakarensis_KOD1 | AP006878 | GCA_000009965,1 | Solfatara on Kodak | 29,22566 | 129,33101 | Solfatara on Kodakara Island | | a1994 | 2.1 | 52 | yes | 91.8 | 818.13 | 96.08 | | | | 2289 | 2245 | 483 | 1231 | 537 | 477 | 54.83296214 | 23.91982183 | 21.24721604 | 21.51447661 |
| Thermococcus_kodakarensis_TS900 | NZ_CP116885,1 | GCA_028471785,1 | culture Solfatara o | 31,596554 | 130,557116 | Solfatara on Kodakara Island | | a1994 | 2.0857 | 52 | yes | 91.03 | 817.67 | 97.85 | | | | 2286 | 2242 | 483 | 1228 | 536 | 478 | 54.77252453 | 23.90722569 | 21.32024978 | 21.54326494 |
| Thermococcus_kodakarensis_TS901 | NZ_CP116886,1 | GCA_028471865,1 | culture Solfatara o | 31,596554 | 130,557116 | Solfatara on Kodakara Island | | a1994 | 2.07848 | 52 | yes | 91.03 | 814.74 | 97.89 | | | | 2282 | 2238 | 483 | 1228 | 535 | 475 | 54.87042002 | 23.90527256 | 21.22430742 | 21.58176944 |
| Palaeococcus_ferrophilus_DSM_13482 | NZ_LANF01000000 | GCA_000966265,1 | Myojin Knoll in the | 32,06 | 139,52 | deep-sea hydrothermal vent chimney | | 1998 | 2.206431 | 54.5 | No | 91.79 | 852.68 | 97.19 | | | | 2293 | 2261 | 483 | 1140 | 471 | 650 | 50.42016807 | 20.83149049 | 28.74834144 | 21.3622291 |
| Thermococcus_thioreducens_OGL_20P | CP015105 | GCA_002214545,1 | Atlantic Ocean: Mi | 36,2 | -33,9 | Marine hydrotherm | 2300 | 1999 | 2.1 | 53.5 | yes | 90.32 | 811.21 | 95.32 | | | | 2287 | 2258 | 483 | 1254 | 718 | 286 | 55.53587245 | 31.79805137 | 12.66607617 | 21.39061116 |
| Pyrococcus_sp_IRI_42C | 8814 | PIRI42C,1 | Mid-Atlantic Ridge | 36,229433 | -33,90369 | deep-sea hydrothe | 2,300m | 2001 | 1.814925 | 41.04 | yes | 90.25 | 799.26 | 112.16 | | | | 2057 | 2013 | 483 | 997 | 434 | 582 | 49.52806756 | 21.5598609 | 28.91207154 | 23.99403875 |
| Thermococcus_sp_IRI29c | 11803 | | Mid Atlantic Ridge | 36,229450 | -33,901967 | hydrothermal chen | 2,274m | 2001 | 2.085848 | 54.54 | yes | 90.98 | 827.54 | 100.07 | | | | 2299 | 2263 | 483 | 1259 | 776 | 228 | 55.63411401 | 34.29076447 | 10.07512152 | 21.34334954 |
| Thermococcus_sp_IRI_05c | 8817 | | Mid Atlantic Ridge | 36,229617 | -33,901382 | 2257 | | 2001 | 2.119293 | 53.14 | yes | 91.81 | 831.38 | 92.41 | 1 | plasmid TIRI05C_p,1 | | 2349 | 2298 | 483 | 1259 | 751 | 288 | 54.78677111 | 32.68059182 | 12.53263708 | 21.01827676 |
| Thermococcus_camini_IRI35c | 11151 | GCA_904067545,1 | Mid-Atlantic Ridge | 36,388089 | -33,858681 | deep-sea hydrothe | 2,250 m | 2001 | 2.022529 | 54.6 | yes | 91.38 | 845.35 | 94.19 | | | | 2166 | 2140 | 483 | 1265 | 710 | 165 | 59.11214953 | 33.17757009 | 7.710280374 | 22.57009346 |
| Thermococcus_IRI_06c | 12654 | | Mid Atlantic Ridge | 36,4266667 | -33,93277778 | | | 2001 | 2.139740 | 55 | yes | 91.65 | 833.79 | 99.31 | 1 | plasmid THIRI06C_p,1 | | 2367 | 2299 | 483 | 1266 | 658 | 375 | 55.06742062 | 28.62113963 | 16.31143976 | 21.00913441 |
| Thermococcus_IRI15c | 11152 | | Mid Atlantic Ridge | 36,4266667 | -33,93277778 | | | 2001 | 2.061457 | 54.6 | yes | 91.46 | 842.13 | 93.62 | | | | 2245 | 2210 | 483 | 1271 | 739 | 200 | 57.51131222 | 33.43891403 | 9.049773756 | 21.85520362 |
| Thermococcus_sp_IRI_33c | 8820 | | Mid Atlantic Ridge, | 36,4266667 | -33,93277778 | Outside part of the | 2,274m | 2001 | 2.115565 | 52.8 | yes | 91.23 | 815.8 | 96.4 | 1 | plasmid TIRI33C_p,1 | | 2376 | 2333 | 483 | 1280 | 767 | 286 | 54.86498071 | 32.87612516 | 12.25889413 | 20.70295757 |
| Thermococcus_sp_IRI10c | 11798 | | Mid Atlantic Ridge | 36,4266667 | -33,93277778 | | | 2001 | 2.124406 | 54.24 | yes | 91.06 | 838.78 | 99.2 | | | | 2314 | 2270 | 483 | 1270 | 755 | 245 | 55.94713656 | 33.25991189 | 10.79295154 | 21.27753304 |
| Thermococcus_sp_IRI14c | 11810 | | Mid Atlantic Ridge | 36,4266667 | -33,93277778 | | | 2001 | 2.064896 | 54.59 | yes | 91.32 | 839 | 96.95 | | | | 2255 | 2221 | 483 | 1271 | 742 | 208 | 57.22647456 | 33.40837461 | 9.365150833 | 21.74696083 |
| Thermococcus_sp_IRI24c | 11800 | | Mid Atlantic Ridge | 36,4266667 | -33,93277778 | | | 2001 | 2.085915 | 54.54 | yes | 91.08 | 828.08 | 98.48 | | | | 2300 | 2264 | 483 | 1259 | 775 | 230 | 55.60954064 | 34.23144876 | 10.1590106 | 21.33392226 |
| Thermococcus_sp_IRI25c | 11809 | | Mid Atlantic Ridge | 36,4266667 | -33,93277778 | | | 2001 | 2.081813 | 54.58 | yes | 91.49 | 840.56 | 95.06 | | | | 2273 | 2236 | 483 | 1273 | 749 | 214 | 56.93202147 | 33.49731664 | 9.570661896 | 21.60107335 |
| Thermococcus_sp_IRI27c2 | 11807 | | Mid Atlantic Ridge | 36,4266667 | -33,93277778 | | | 2001 | 2.085915 | 54.54 | yes | 90.95 | 832.16 | 100.01 | | | | 2285 | 2247 | 483 | 1258 | 777 | 212 | 55.98575879 | 34.57943925 | 9.434801958 | 21.4953271 |
| Thermococcus_sp_P6 | CP015104 | GCA_002214525,1 | Greece, Palaeocho | 36,58 | 24,38 | hydrothermal sedi | 10 | 1997 | 1.52282 | 54.9 | yes | 90.89 | 857.87 | 87.39 | | | | 1599 | 1589 | 483 | 1134 | 286 | 169 | 71.36563877 | 17.99874135 | 10.63561989 | 30.39647577 |
| Thermococcus_aegaeus_DSM_12767 | 11780 | THAEA,1 | Aegean See, Palae | 36,62 | 24,44 | marine hydrotherm | 9,5m | 1996 | 1.915151 | 42.96 | yes | 91.42 | 843.57 | 96.02 | | | | 2085 | 2030 | 483 | 1121 | 436 | 473 | 55.22167488 | 21.47783251 | 23.30049261 | 23.79310345 |

| | | | | | | | | | | | | | | | | | | | | | | | | | | | |
|---|---|---|---|---|---|---|---|---|---|---|---|---|---|---|---|---|---|---|---|---|---|---|---|---|---|---|---|
| Pyrococcus_furiosus_DSM_3638 | AE009950 | GCA_000007305,1 | Vulcano Island Italy | 38,396 | 14,968 | Geothermally heated marine sedimen | | a1982 | 1.90826 | 40.8 | yes | 91.87 | 808.71 | 100.35 | | | | 2027 | 1957 | 483 | 1010 | 498 | 449 | 51.60960654 | 25.44711293 | 22.94328053 | 24.68063362 |
| Pyrococcus_furiosus_COM1 | CP003685 | GCA_000275605,1 | Vulcano Island Italy | 38,396 | 14,968 | Geothermally heated marine sedimen | | a1982 | 1.909827 | 40.8 | yes | 91.37 | 804.19 | 103.8 | | | | 2033 | 1961 | 483 | 1009 | 500 | 452 | 51.45334013 | 25.49719531 | 23.04946456 | 24.63029067 |
| Pyrococcus_furiosus_DSM_3638_Reg | | GCA_008245085 | pure culture of Pyr | 38,396 | 14,968 | Geothermally heated marine sedimen | | a1982 | 1.889914 | 40.75 | yes | 90 | 822.16 | 102.31 | | | | 2018 | 1962 | 483 | 1017 | 494 | 451 | 51.83486239 | 25.1783894 | 22.98674822 | 24.617737 |
| Thermococcus_celer_Vu_13 | CP014854 | GCA_002214365,1 | Italy: volcano islan | 38,399179 | 14,9693 | Marine hydrothermal vent | | a1982 | 1.9 | 56.5 | yes | 89.42 | 821.51 | 95.22 | | | | 1961 | 1927 | 483 | 1227 | 465 | 235 | 63.67410483 | 24.13077322 | 12.19512195 | 25.06486767 |
| Thermococcus_alcaliphilus_DSM_10322 | 11782 | THALCA,1 | Porto di Levante, V | 38,416205 | 14,331 | a shallow beach | 1 m | <1995 | 1.891997 | 43.03 | yes | 91.85 | 840.92 | 90.24 | | | | 2076 | 2043 | 483 | 1131 | 465 | 447 | 55.35976505 | 22.76064611 | 21.87958884 | 23.64170338 |
| Palaeococcus_helgesoni_DSM_15127 | 11779 | PHEL,1 | geothermal well, V | 38,417610 | 14,957505 | a shallow geothermal well that taps m | | 1999 | 1.953937 | 47.01 | yes | 90.5 | 836.39 | 108.41 | | | | 2126 | 2105 | 483 | 1066 | 435 | 604 | 50.64133017 | 20.66508314 | 28.6935867 | 22.94536817 |
| Thermococcus_celer_SH1 | | | Black Point, 'île de | 38,6219549288500 | 15,13240287 | hydrothermal chim | 20 | 2017 | 1.890105 | 56.5 | yes | 90.94 | 800.9 | 98.68 | | | | 2014 | 1975 | 483 | 1232 | 482 | 261 | 62.37974684 | 24.40506329 | 13.21518987 | 24.4556962 |
| Thermococcus_litoralis_DSM_5473 | CP006670 | GCA_000246985,3 | Naples, Bay of Luc | 40,80 | 14,1 | shallow submarine | shallow | 1985 | 2.19183 | 43.1 | yes | 90.78 | 777.97 | 98.6 | | | | 2387 | 2295 | 483 | 1101 | 568 | 626 | 47.97385621 | 24.74945534 | 27.27668845 | 21.04575163 |
| Thermococcus_sp_2319x1 | CP012200 | GCA_001484685,1 | Kunashir Island, Ru | 44,188 | 145,8411 | hot vent located ir | shallow | 2012 | 1.96122 | 44.6 | yes | 90.92 | 758.43 | 98.56 | | | | 2082 | 2024 | 483 | 1103 | 454 | 467 | 54.49604743 | 22.43083004 | 23.07312253 | 23.86363636 |
| Thermococcus_sp_2319x1_Essen | | GCA_902813195 | Kunashir Island, Ru | 44,188 | 145,8411 | hot vent located in the tidal zone of K | | 2012 | 1.989851 | 44.39 | yes | 88.23 | 802.46 | 108.6 | | | | 2130 | 2077 | 483 | 1102 | 472 | 503 | 53.05729417 | 22.72508426 | 24.21762157 | 23.25469427 |
| Thermococcus_cleftensis_CL1 | CP003651 | GCA_000265525,1 | north-eastern Paci | 46,026460 | -130,05679 | polychaete worm c | 2,350m | 1994 | 1.95031 | 55.8 | yes | 90.98 | 837.17 | 95.62 | | | | 2063 | 2040 | 483 | 1253 | 575 | 212 | 61.42156863 | 28.18627451 | 10.39215686 | 23.67647059 |
| Pyrococcus_sp_ST04 | CP003534 | GCA_000263735,1 | Endeavour Segmen | 46,160035 | -129,989019 | deep-sea hydrothe | 2,290m | 2004 | 1.73689 | 42.3 | yes | 91.42 | 874.02 | 94.34 | | | | 1809 | 1792 | 483 | 1023 | 485 | 284 | 57.08705357 | 27.06473214 | 15.84821429 | 26.953125 |
| Thermococcus_barossii_SHCK_94 | CP015101 | GCA_002214465,1 | Pacific Ocean, Jua | 46,160035 | -129,989019 | Hydrothermal vent | 2,100m | a1998 | 1.9 | 54.5 | yes | 91.81 | 863.63 | 86.47 | | | | 2030 | 2014 | 483 | 1274 | 561 | 179 | 63.2571996 | 27.8550149 | 8.887785501 | 23.98212512 |
| Thermococcus_paralvinellae_ES1 | CP006965 | GCA_000517445,1 | Endeavour Segmen | 47,57 | -129,06 | isolated from a pol | 2200 | 1987 | 2 | 40.5 | yes | 91.5 | 828.18 | 95.64 | | | | 2105 | 2059 | 483 | 1110 | 461 | 488 | 53.90966489 | 22.38950947 | 23.70082564 | 23.45798932 |
| Thermococcus_stetteri_DSM_5262 | 11785 | | Northern Kurils, Us | 47,570 | 152,82 | marine volcanic crater fields of Krater | | a1989 | 2.03478 | 52.5 | yes | 89.69 | 790.06 | 110.85 | | | | 2326 | 2216 | 483 | 1251 | 499 | 466 | 56.45306859 | 22.51805054 | 21.02888087 | 21.79602888 |
| Pyrococcus_endeavori_ES4 | 4064 | PYES4,1 | Endeavour Segmen | 47,95 | -129,05 | hydrothermal chim | 2,000m | | 1.710181 | 42.15 | yes | 92.63 | 869.04 | 90.89 | | | | 1829 | 1812 | 483 | 1020 | 461 | 331 | 56.29139073 | 25.4415011 | 18.26710817 | 26.65562914 |
| Thermococcus_bergensis_T7324 | SAMN13612049 | GCA_020386975,1 | Norway: Deep and | 56,3666667 | 3,266666389 | oil field waters | | 1993 | 2.077832 | 44 | no | 90.65 | 770.22 | 96.76 | | | | 2267 | 1944 | 483 | 1076 | 450 | 418 | 55.34979424 | 23.14814815 | 21.50205761 | 24.84567901 |
| Thermococcus_sibiricus_MM_739 | CP001463 | GCA_000022545,1 | Western Siberia, R | 61,116 | 76,75 | formation water, h | 2,350m | 1998 | 1.845800 | 40.2 | yes | 90.57 | 799.1 | 91.14 | | | | 1970 | 1928 | 483 | 1068 | 396 | 464 | 55.39419087 | 20.53941909 | 24.06639004 | 25.05186722 |
| Thermococcus_sp_CIR_03a | 8822 | TCIR03A,1 | Rodrigues triple ju | 70 | 25 | | 2420 | 2001 | 2.207808 | 53.55 | yes | 92.24 | 852.49 | 91.05 | | | | 2397 | 2357 | 483 | 1276 | 709 | 372 | 54.13661434 | 30.08061095 | 15.78277471 | 20.49215104 |
| Thermococcus_sp_CIR_10a | 8818 | TCIR10A,1 | Rodrigues triple ju | 70 | 25 | | 2420 | 2001 | 2.201239 | 41.38 | yes | 92.02 | 812.92 | 90.36 | | | | 2510 | 2444 | 483 | 1144 | 562 | 738 | 46.80851064 | 22.99509002 | 30.19639935 | 19.76268412 |
| Thermococcus_henrietii_EXT12c | LT900021,1 | GCA_900198835,1 | East Pacific Ridge | 9,844500 | -104,293833 | deep-sea hydrothe | 2,496m | 2001 | 2.15576 | 54.6 | yes | 91.05 | 836.54 | 101.66 | | | | 2325 | 2287 | 483 | 1241 | 723 | 323 | 54.26322693 | 31.61346742 | 14.12330564 | 21.11937035 |
| Pyrococcus_sp_EXT_15c | 8813 | PEXT15C,1 | East Pacific Ridge | 9,844500 | -104,293833 | | | 2001 | 1.851203 | 45.01 | yes | 92.5 | 816.61 | 88.22 | | | | 2111 | 2064 | 483 | 1048 | 524 | 492 | 50.7751938 | 25.3875969 | 23.8372093 | 23.40116279 |
| Pyrococcus_EXT16 | 7485 | PYEXT16,1 | East Pacific Ridge | 9,844500 | -104,293833 | | | 2001 | 1.952677 | 45.05 | yes | 91.95 | 811.13 | 97.54 | 1 | pPYEXT16 | PYEXT16_p,1 | 2230 | 2168 | 483 | 1045 | 620 | 503 | 48.20110701 | 28.59778598 | 23.20110701 | 22.27859779 |
| Thermococcus_sp_E10P11 | 11811 | TE10P11,1 | East Pacific Ridge | 9,844500 | -104,293833 | | | 2001 | 1.963818 | 54.95 | yes | 91.89 | 834.25 | 97.44 | | | | 2171 | 2142 | 483 | 1242 | 770 | 130 | 57.98319328 | 35.94771242 | 6.069094304 | 22.54901961 |
| Thermococcus_sp_E10P8 | 11801 | TE10P8,1 | East Pacific Ridge | 9,844500 | -104,293833 | | | 2001 | 1.963818 | 54.95 | yes | 91.88 | 834.56 | 97.55 | | | | 2170 | 2141 | 483 | 1241 | 770 | 130 | 57.96356843 | 35.96450257 | 6.071929005 | 22.55955161 |
| Thermococcus_sp_E15p33 | 8816 | TE15P33,1 | East Pacific Ridge | 9,844500 | -104,293833 | | | 2001 | 2.048119 | 54.77 | yes | 92.3 | 826.95 | 91.16 | | | | 2294 | 2268 | 483 | 1248 | 775 | 245 | 55.02645503 | 34.17107584 | 10.80246914 | 21.2962963 |
| Thermococcus_E15P35 | 10971 | | East Pacific Ridge | 9,844500 | -104,293833 | hydrothermal source | | 2001 | 2.049601 | 54.77 | yes | 92.41 | 825.05 | 89.5 | 1 | plasmid TE15P35,2 | | 2317 | 2287 | 483 | 1248 | 775 | 264 | 54.56930477 | 33.88718846 | 11.54350678 | 21.11937035 |
| Thermococcus_EXT11c | 11137 | | East Pacific Ridge | 9,844500 | -104,293833 | | | 2001 | 1.909018 | 54.95 | yes | 92.25 | 825.54 | 88.57 | | | | 2140 | 2110 | 483 | 1239 | 646 | 225 | 58.72037915 | 30.61611374 | 10.66350711 | 22.89099526 |
| Thermococcus_sp_EXT09c | 8819 | | East Pacific Ridge | 9,844500 | -104,293833 | Outside part of the cheminey | | 2001 | 1.922260 | 54.82 | yes | 92.2 | 814.8 | 86.03 | 2 | plasmid TEXT09C_pb,1; plasmid TEXT( | | 2183 | 2147 | 483 | 1237 | 675 | 235 | 57.61527713 | 31.43921751 | 10.94550536 | 22.49650675 |
| Thermococcus_sp_DS1 | 7110 | Chromosome,1 | Pacific Ocean:East-Pacific Rise | | | | | | 2.135121 | 52.66 | yes | 91.17 | 811.71 | 97.29 | | | | 2405 | 2349 | 483 | 1277 | 765 | 307 | 54.36355896 | 32.56704981 | 13.06939123 | 20.56194125 |
| Thermococcus_sp_unknown | 8821 | | unknown | | | | | | 1.924085 | 54.73 | yes | 92.07 | 829.42 | 90.3 | 1 | plasmid THERMO_p,1 | | 2141 | 2119 | 483 | 1244 | 661 | 214 | 58.70693723 | 31.19395941 | 10.09910335 | 22.79377065 |
