## Supplementary figures and images for "Thermococcales pangenome uncovers new genera and modular evolution"

### Supplementary figure 1

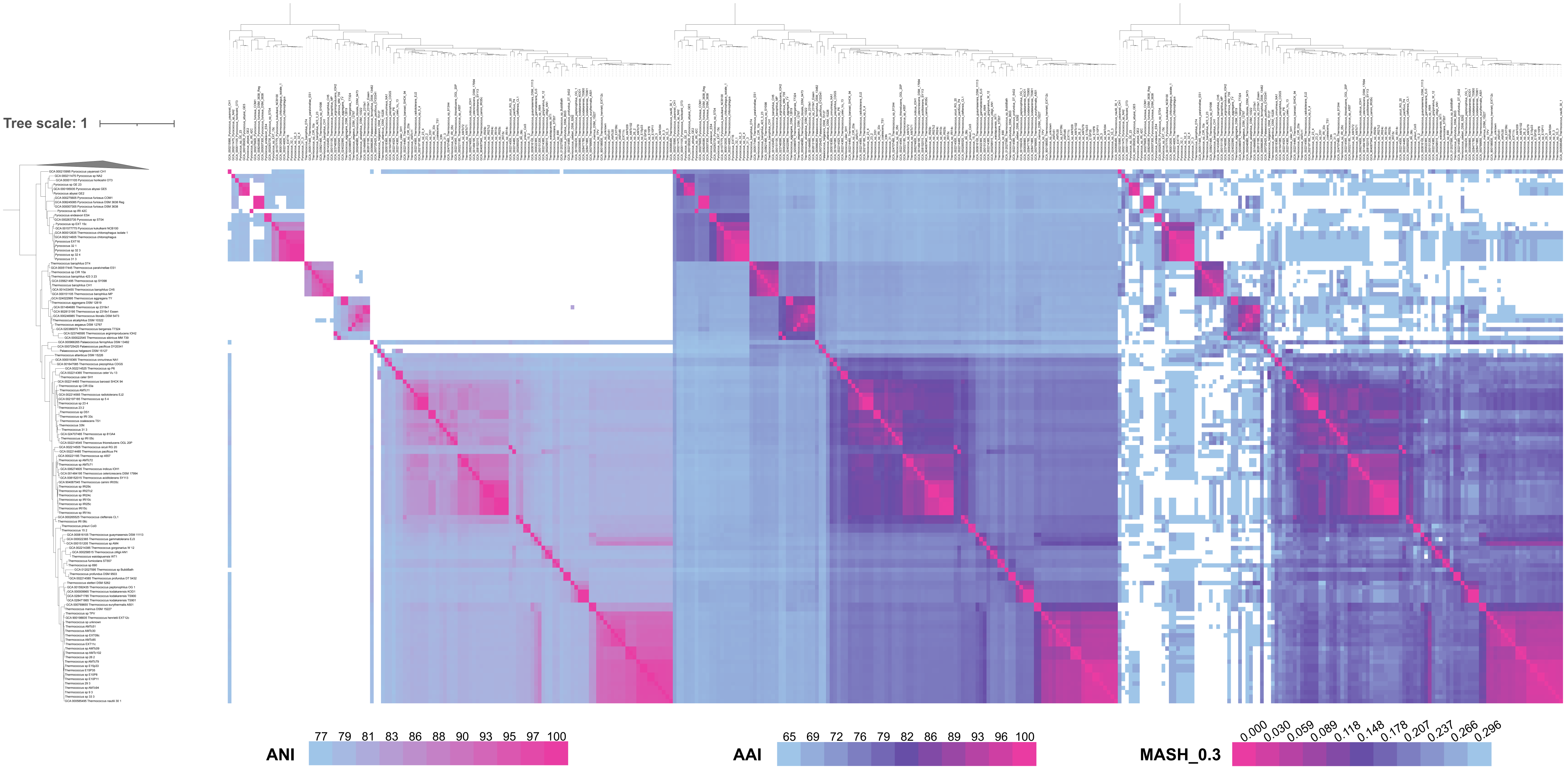

### Supplementary figure 2

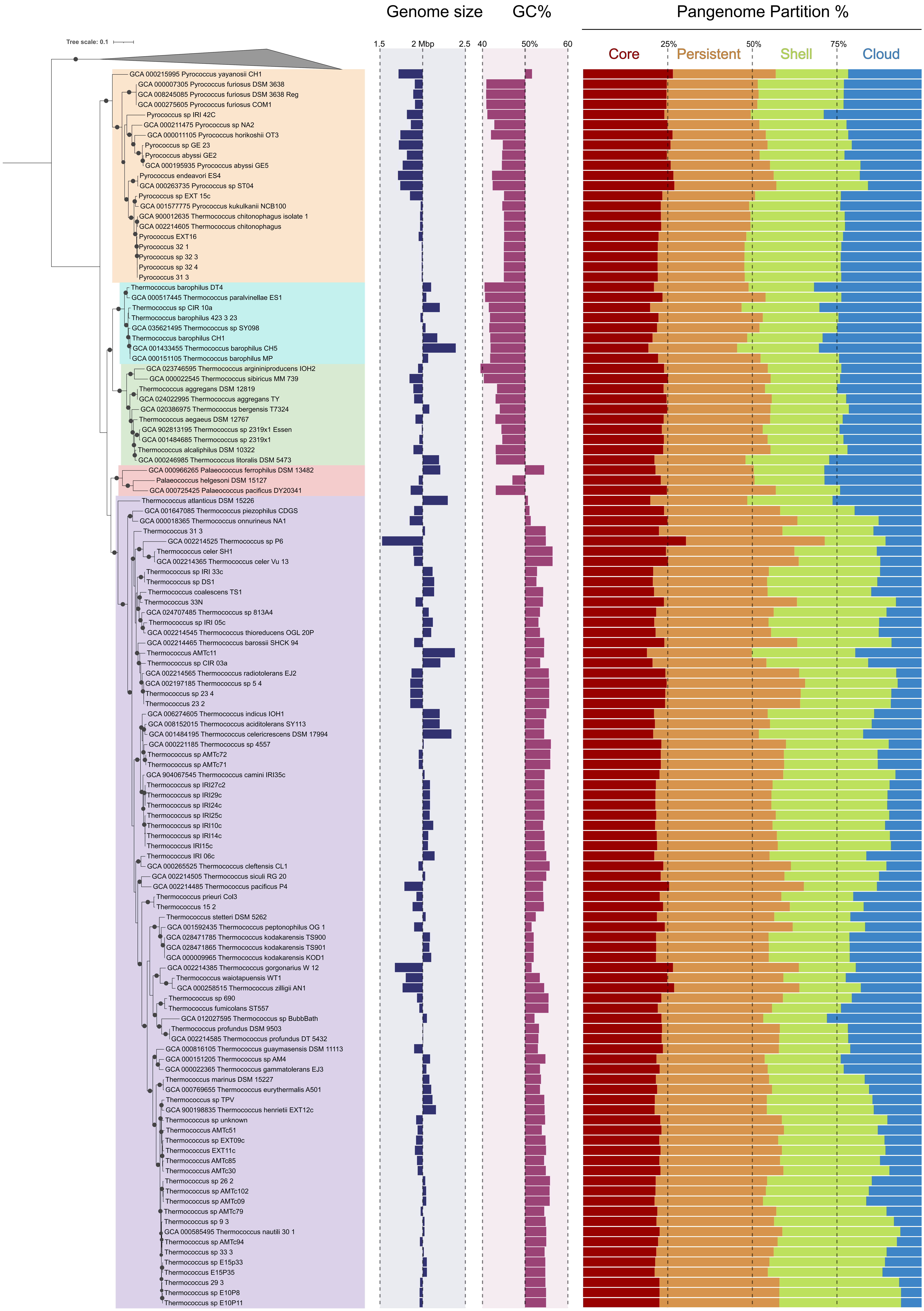

### Supplementary figure 3

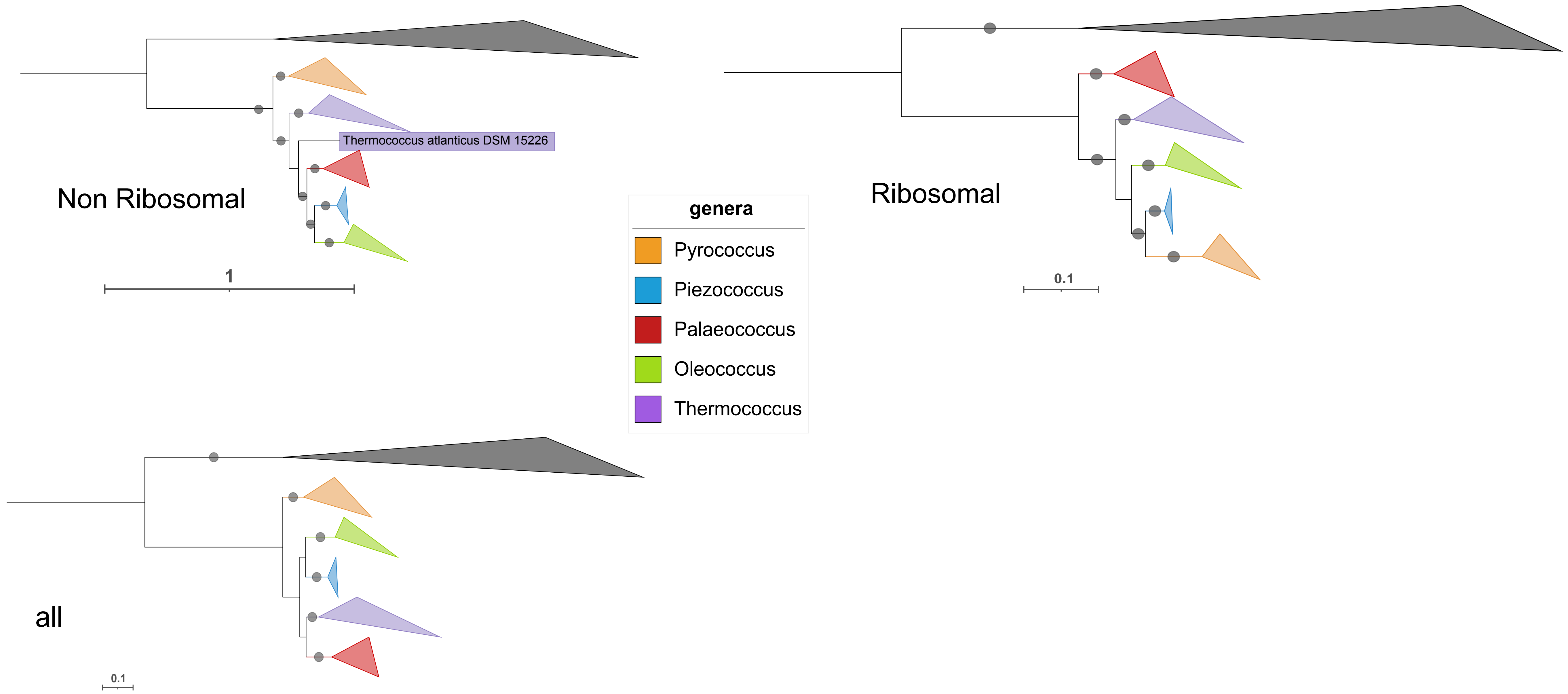

### Supplementary figure 4

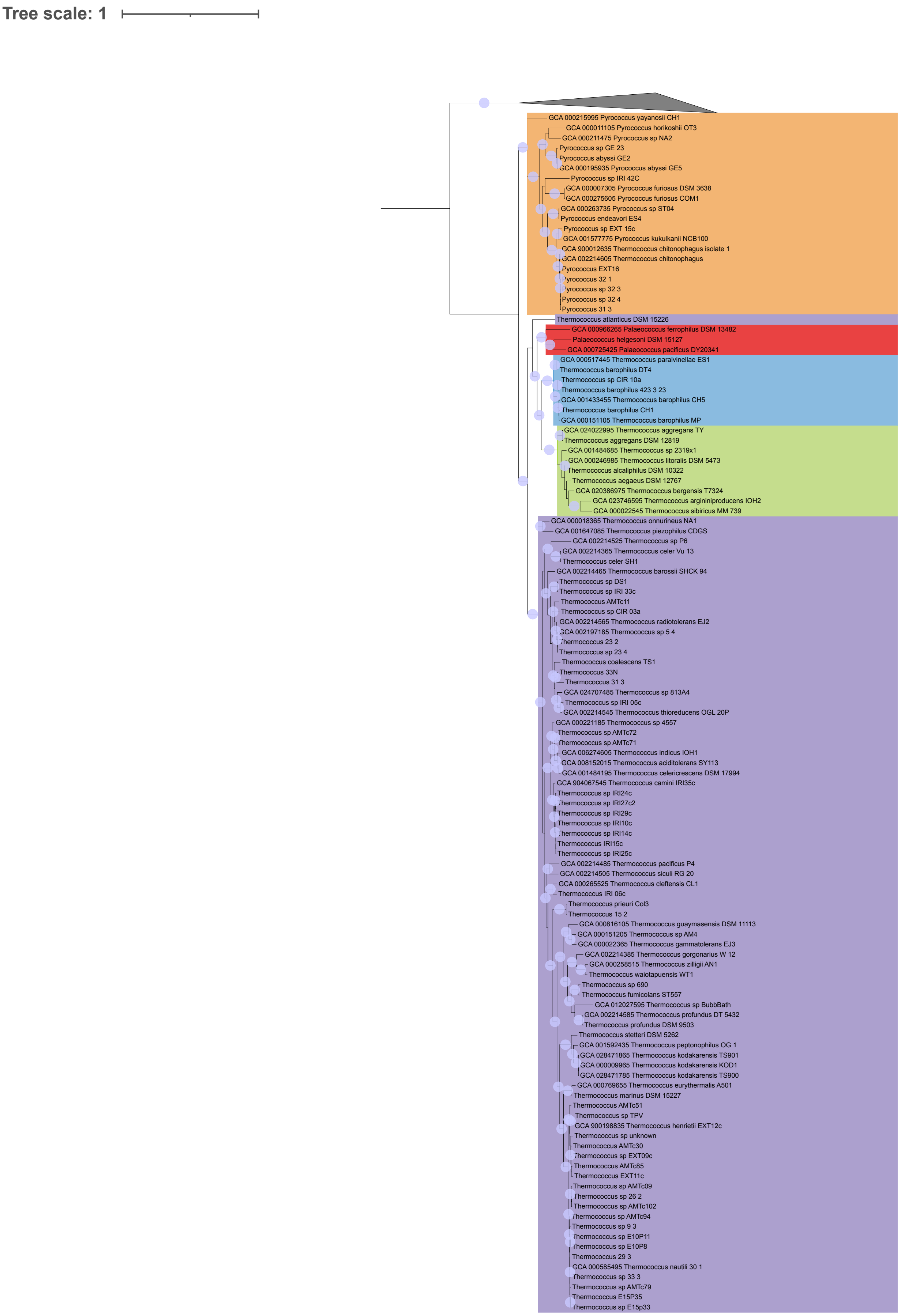

### Supplementary figure 5

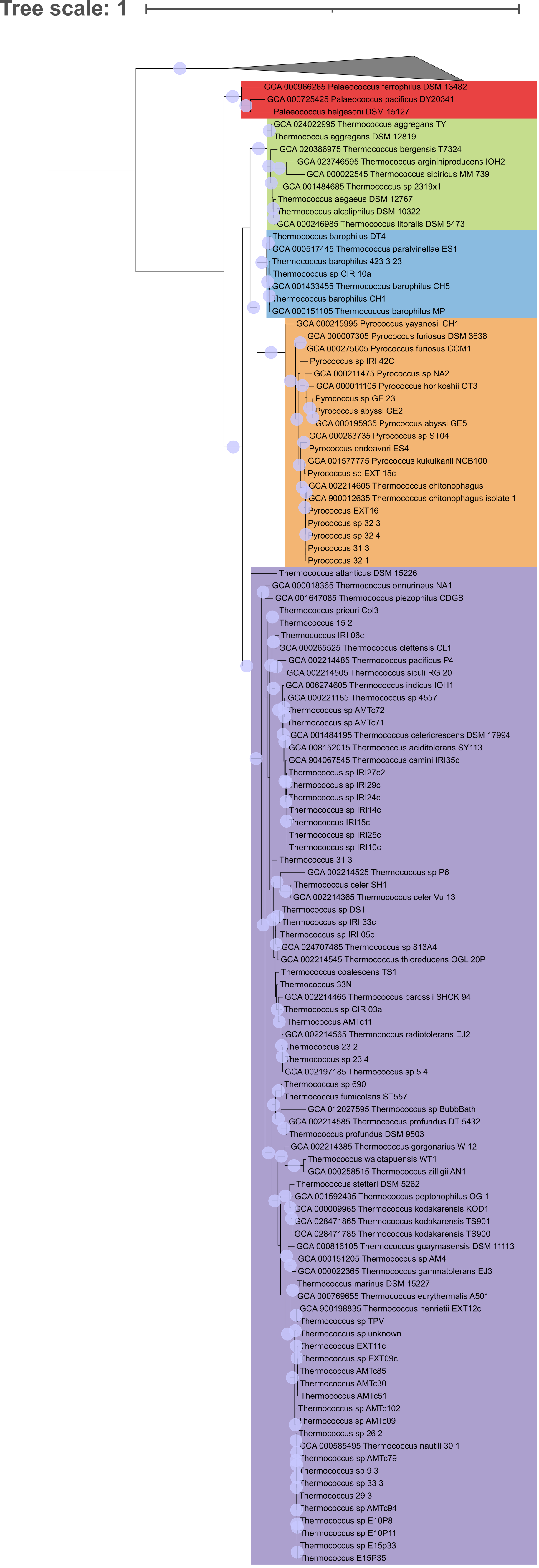

### Supplementary figure 6

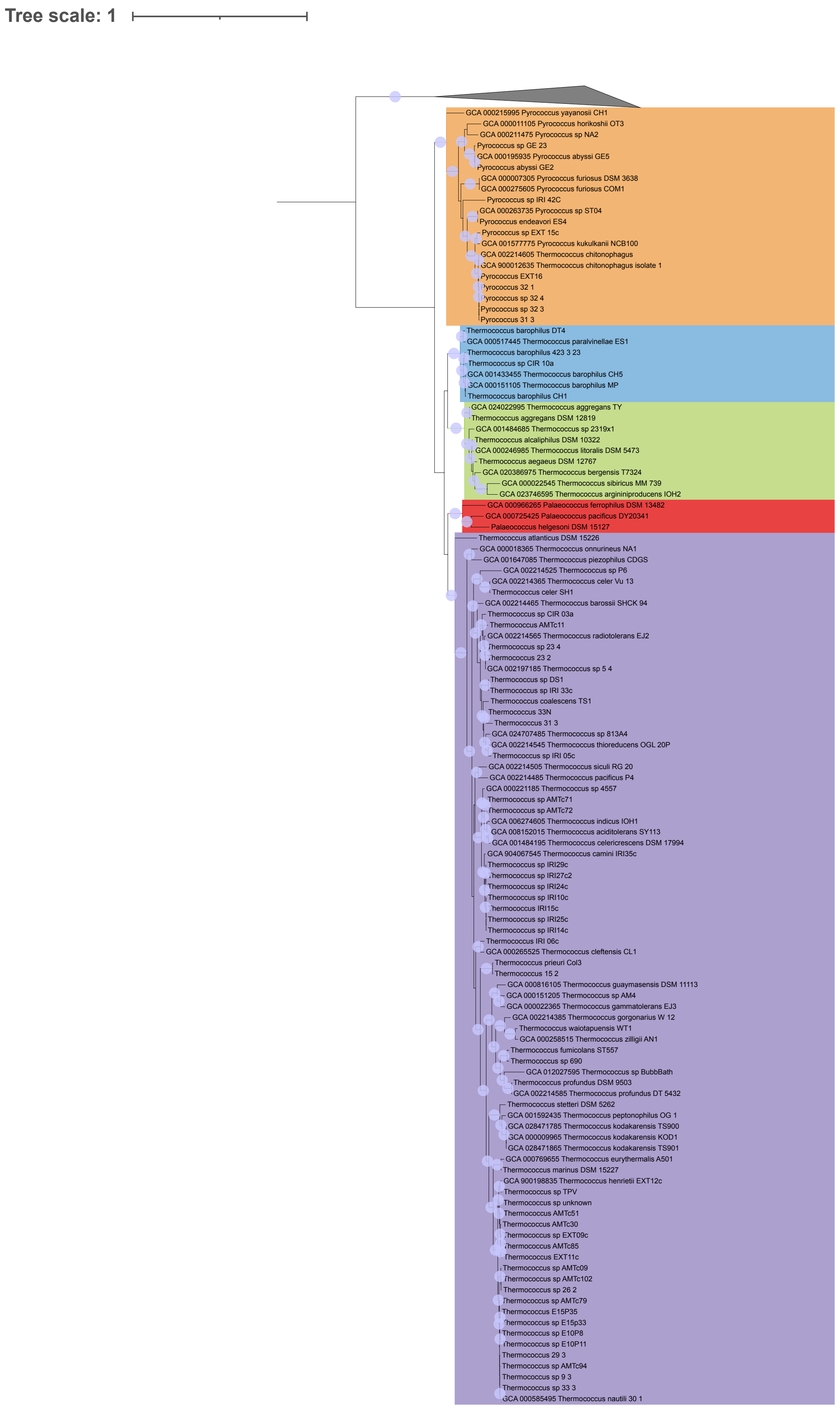

### Supplementary figure 7

#### 32971837 = HypE

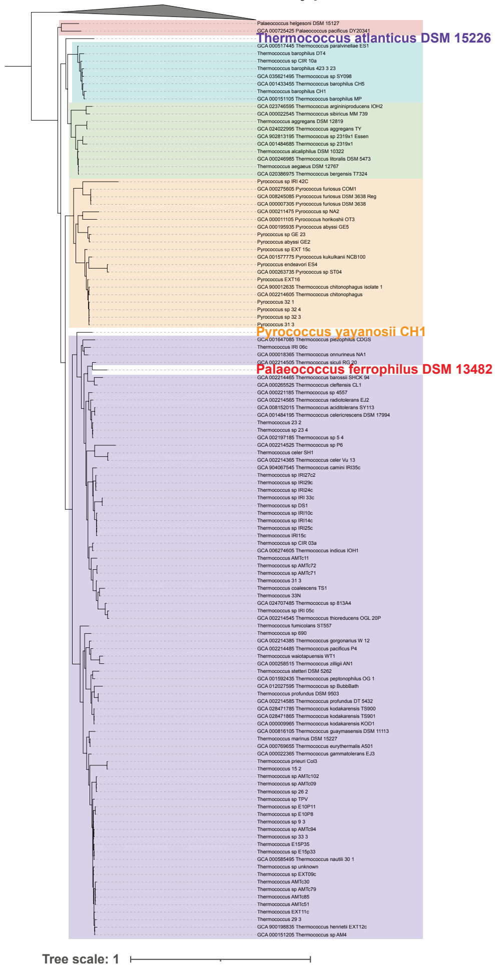


#### 32979154 = slipin family protein

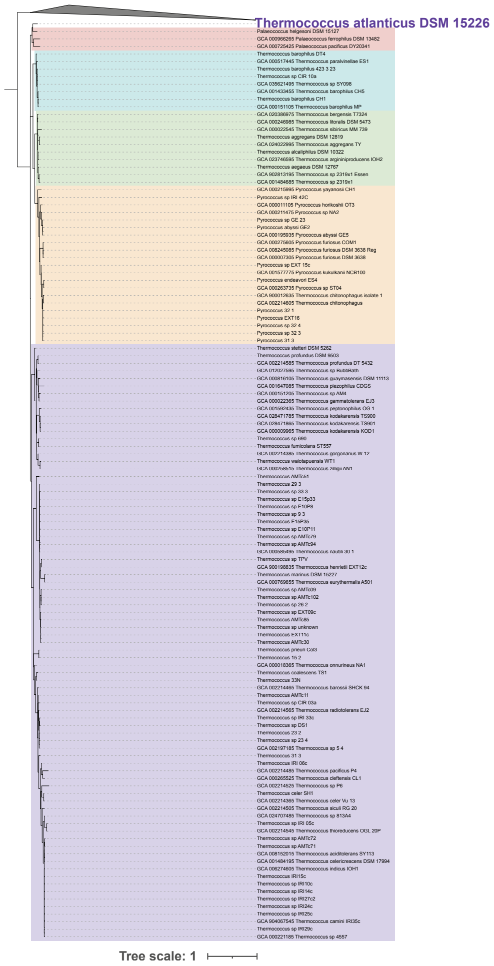


#### 32980655 = HypF

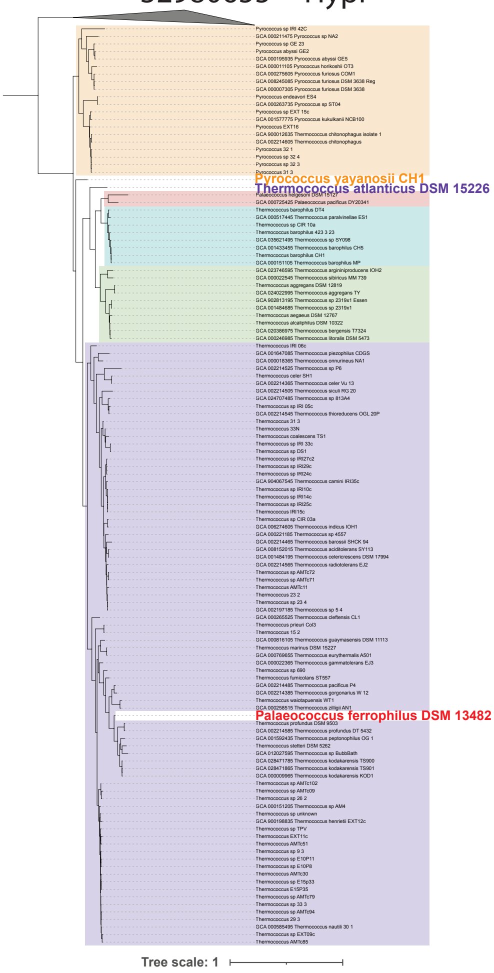


#### 52625396 = UbiD family decarboxylase

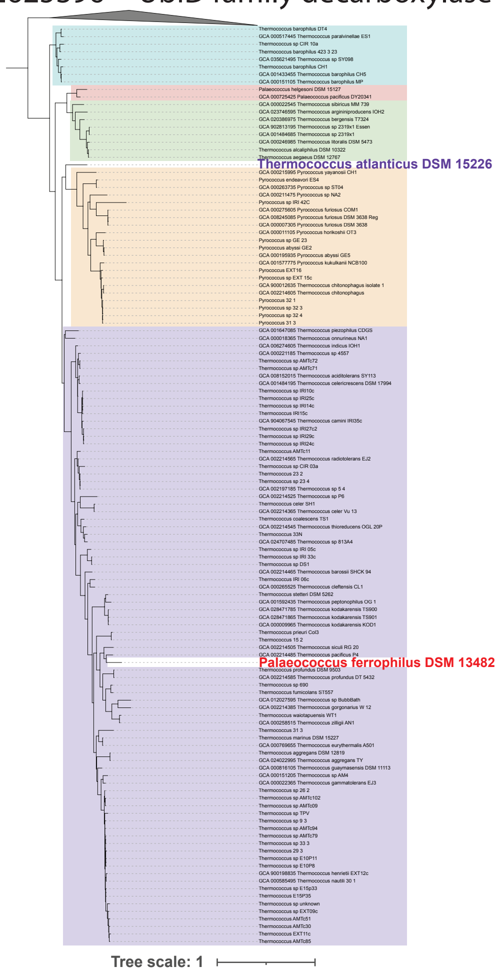

### Supplementary figure 8

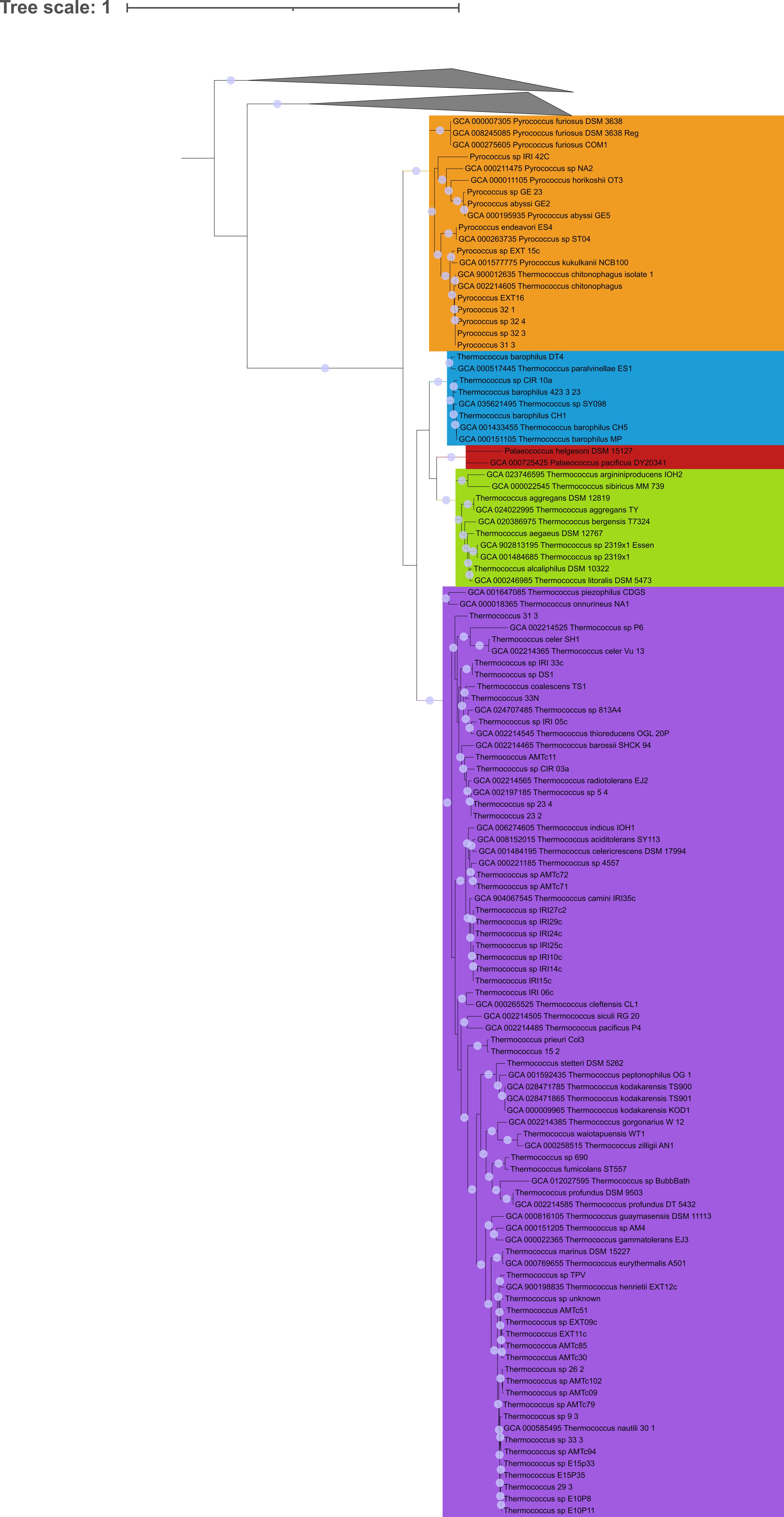

### Supplementary figure 9

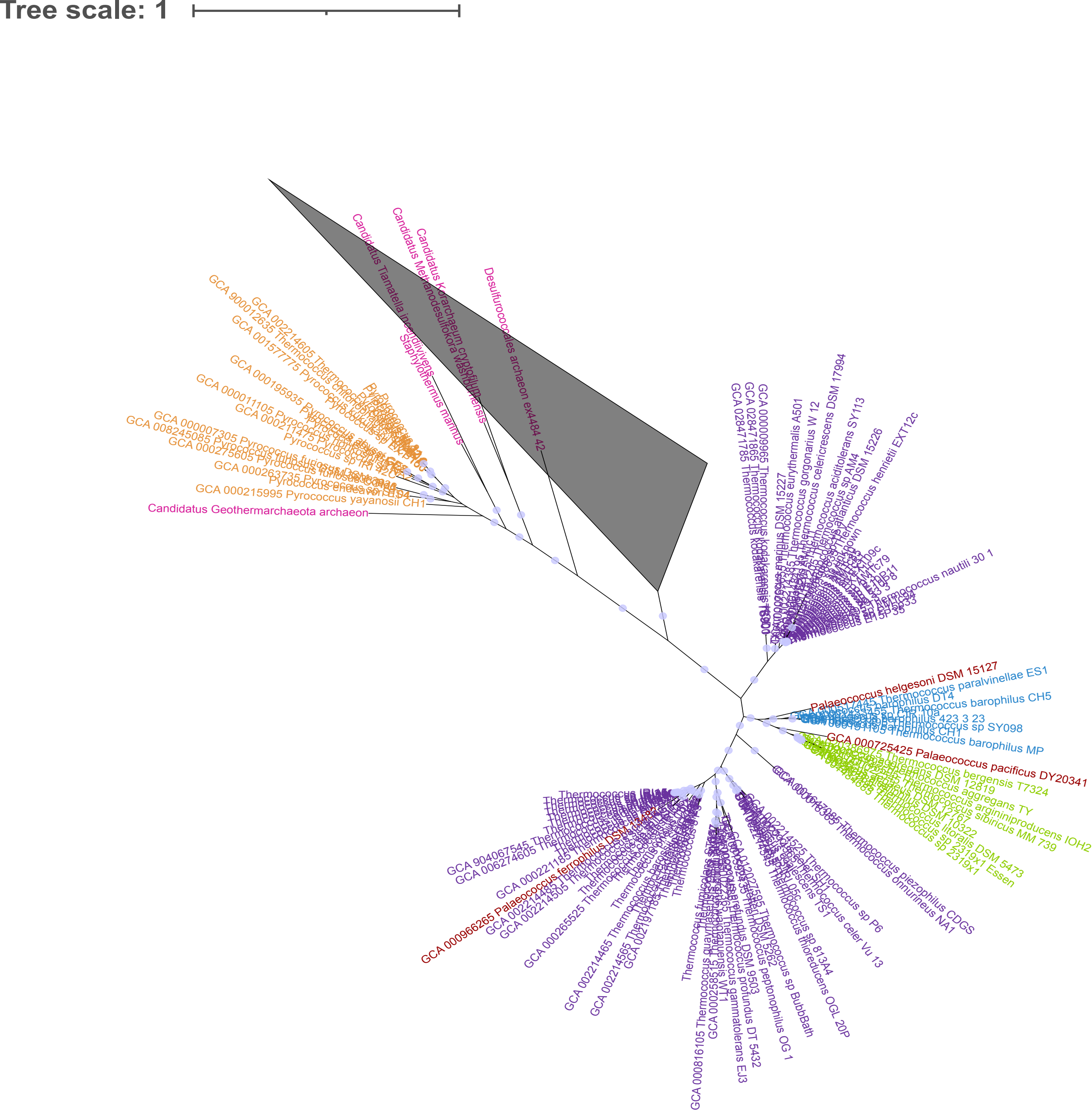
